# Quantifying potential abiotic drivers of the ‘nurse-plant effect’ in two dominant shrub species of the northern Chihuahuan Desert

**DOI:** 10.1101/2023.11.16.567462

**Authors:** Scott Ferrenberg, Akasha M. Faist, Brooke Osborne, Steven R. Lee

## Abstract

Aggregations of plants surrounded by areas without vegetation cover in dryland ecosystems are thought to arise when larger plants facilitate the recruitment and/or performance of smaller “protégé” plants—a phenomenon referred to as the “nurse-plant” effect. While numerous drivers can generate a nurse-plant effect, efforts to quantify multiple drivers simultaneously are rare. After verifying a higher density of protégés beneath the foundational shrubs *Larrea tridentata* and *Prosopis glandulosa*, multiple potential mechanisms underlying the nurse-plant effect were quantified in the Chihuahuan Desert of southern New Mexico. Comparisons of properties under shrub canopies relative to unvegetated interspaces revealed significantly greater concentrations of soil nutrients and lower photosynthetically active radiation and soil temperatures beneath shrubs but a consistently higher soil moisture in interspaces despite a greater water holding capacity in soils beneath shrubs. Nutrient concentrations were greater, on average, in soils beneath *P. glandulosa* than *L. tridentata* but protégé plant numbers did not significantly differ among the species. Further, in *L. tridentata* and *P. glandulosa,* canopy size was positively related to levels of understory shading, and canopy size of *P. glandulosa* was also positively related to soil nitrogen and microbial biomass. Results of this study suggest that a majority of the variance in the abiotic nurse-plant effect of this low-latitude system is explained by radiation interception and the concomitant reduction in temperatures experienced by protégé plants as opposed to direct effects of shrubs on soil water availability. As global change pressures intensify in drylands, a loss of perennial plant cover through mortality or dieback in canopies could have substantial, negative effects on soil biogeochemical pools and plant diversity. Additional quantification of the spatial and temporal variance in different mechanisms driving the nurse-plant effect across environmental and climatic gradients is needed to improve our understanding of plant community dynamics in dryland ecosystems.

## 1. Introduction

Plant aggregations resulting from facilitation have long been argued to be common in dryland ecosystems (Flores & Jurado, 2003; Maestre et al., 2009). In cases where a larger perennial plant modifies the understory environment to facilitate the recruitment, growth, or survival of others, this phenomenon is described as a “nurse-plant” effect (Filazzola & Lortie, 2014). This nomenclature arises from the view that the nurse plant creates a “nursery” where stresses are reduced and conditions favor “protégé plants” that otherwise would not recruit or persist through key life stages (Franco & Nobel, 1989). The original, and most frequently invoked, mechanism driving the nurse-plant effect is a reduction in water stress experienced by protégé plants driven by the reduction in solar radiation beneath the nurse plant’s canopy (Filazzola & Lortie, 2014). However, plants can also improve protégé performance by modifying the movement of seeds and soil particles and increase concentrations of litter and soil nutrients under their canopies (Schlesinger et al., 1996). The importance of soil modification by larger perennial plants for structuring vegetation cover is predicted to increase as a result of desertification processes. Specifically, a loss of grass cover due to the combined pressures of drought and non-native livestock grazing is increasing shrub cover as well as the size and extent of unvegetated soil interspaces in many semi-arid ecosystems (Hoover et al., 2020). Resulting erosion leads to a loss of soil and nutrients from expanding interspaces, leaving the soil retained and deposited around surviving shrubs to be comparatively higher in nutrients (Young et al., 2021). This process leads to spatially discrete patches centered around shrubs that are referred to as “islands of fertility” (Peterjohn & Schlesinger, 1990; Schlesinger et al., 1996) which can promote plant aggregations that reflect variation in soil resource pools.

While it is clear that a combination of above- and belowground drivers can influence the likelihood and degree of plant aggregations in drylands, efforts to isolate and quantify the effects of multiple drivers individually and simultaneously remain rare. Furthermore, like all ecological processes, nurse-plant effects are almost certain to vary across spatial gradients, to change in strength temporally within and among locations, and to vary as a function of shrub species, size, and architecture given the effect of these attributes on shading intensity, litter inputs, and rooting characteristics among other factors (Filazzola & Lortie, 2014). For example, interactions among plants are argued to range from competitive to facilitative as a non-linear, parabolic function of physiological “stress” caused by resource availability (Maestre et al., 2009). Thus, where conditions favor positive, biotic interactions—i.e., facilitation—plant aggregations should arise and be common. Determining which drivers dictate the balance and variability in nurse-plant effects on protégés and to rank their importance across systems is needed if we are to improve our understanding of how changes in climate, land-use, and plant covers will affect dryland functioning.

Prior work in the Chihuahuan Desert has established a positive association among shrubs and understory protégé plants (Badano et al., 2016; Kidron & Gutschick, 2013; Pérez-Sánchez et al., 2015). Using paired surveys beneath shrubs and neighboring interspaces, we verified that fewer annual plants and perennial seedlings occurred in interspaces compared to beneath the canopies of two widespread, dominant shrubs—*Larrea tridentata* (creosote bush) and *Prosopis glandulosa* (honey mesquite)—throughout the summer monsoon growing seasons of 2020 and 2021 (Ferrenberg et al., unpublished data). We then quantified multiple potential mechanisms underlying the observed nurse-plant effect, including measures of solar radiation and soil temperature/moisture beneath shrubs versus interspaces. *Larrea tridentata* is evergreen while *P. glandulosa* is deciduous and also hosts nitrogen (N) fixing bacteria in its roots. Our goal was to compare soil biogeochemical properties, soil temperature and moisture, and incoming photosynthetically active radiation (PAR) under shrub canopies (as well as under shade-cloth ‘canopies’ used as shrub-surrogates in the case of soil moisture; see methods for soil moisture) relative to neighboring unvegetated interspaces to quantify each as a potential driver of the nurse plant effect we observed (Figure 1). We hypothesized that:

1. Soils beneath shrub canopies would have higher carbon and nutrient concentrations compared to interspaces and soils would be less fertile under *L. tridentata* than under the N-fixing *P. glandulosa*.
2. Shrub influences on radiation and soil microclimate would differ as a function of shrub size; we expected shading to increase and soil temperature to decrease as shrub size/cover increased.
3. Shading/reduction in soil temperatures would increase available soil moisture by prolonging soil water volume after precipitation events under shrubs relative to interspaces.

**Figure 1.**
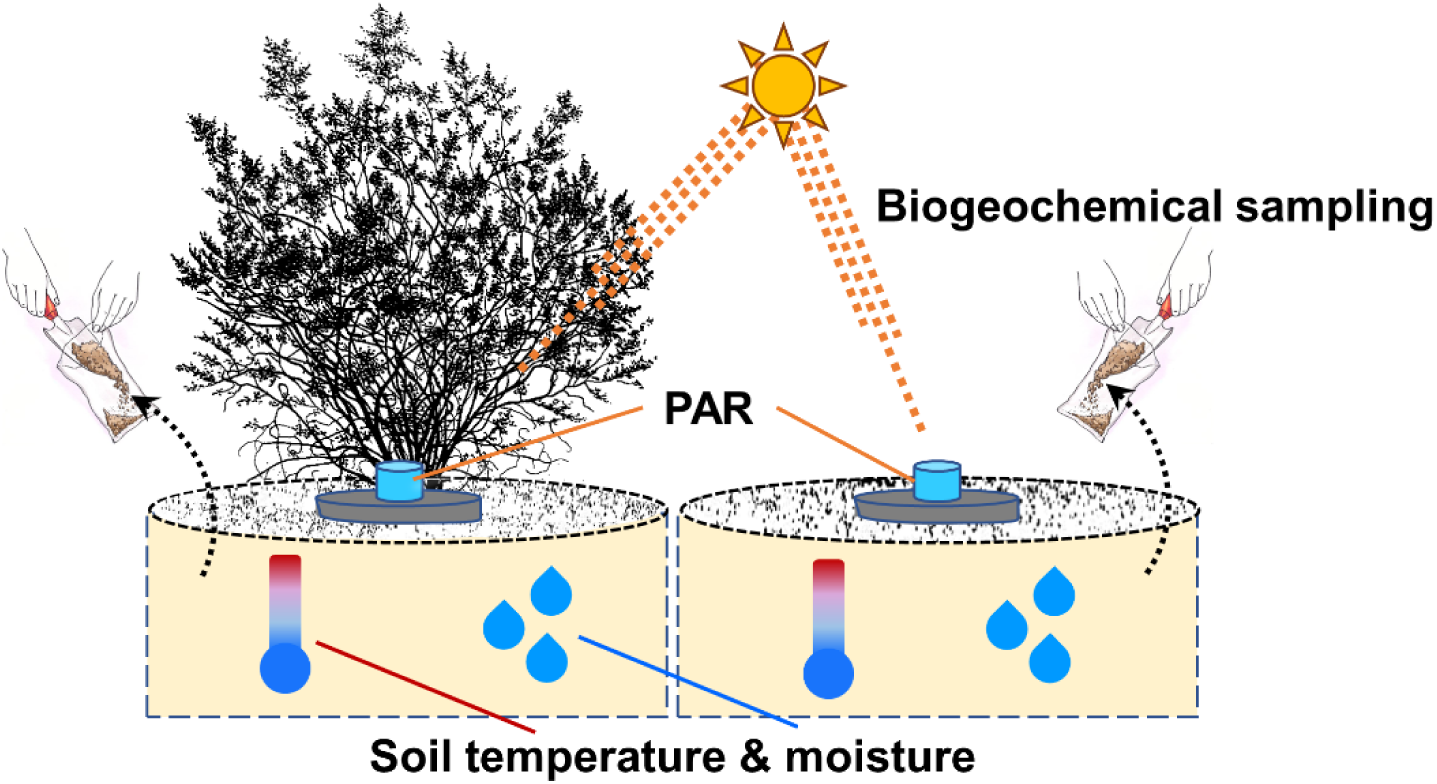
Conceptual design for quantifying shrub effects on incoming photosynthetically active radiation (PAR), soil biogeochemistry, and soil temperature and moisture. Shrubs of *Larrea tridentata* (evergreen) and *Prosopis glandulosa* (deciduous N-fixer) were paired with nearby, unvegetated interspaces. Differences between interspaces and beneath shrub canopies (as well as beneath shade-cloth canopy surrogates for soil moisture) were used to quantify the shrub effect.

## 2. Methods

### 2.1 Study site

We conducted this study between June 2020 and October 2021 at New Mexico State University’s Chihuahuan Desert Rangeland Research Center in southern New Mexico, USA (Figure S1a). The study site was roughly a 300 m × 300 m polygon, bisected by an access road, centered at 32.504558, −106.787015. The climate is semiarid with mean annual temperature of 15.9 °C and mean annual precipitation of 265.7 mm over the century prior to our study (PRISM Climate Group, 2022). Most precipitation falls as a summer monsoon that coincides with the highest daily temperatures each year (Figure S2). Study blocks and plots were located between 1371 and 1378 m above sea level on a piedmont fan roughly 1.6 km east of the base of the Doña Ana Mountains. Our study focused on the evergreen shrub *Larrea tridentata* (creosote bush) and the deciduous, nitrogen-fixing shrub *Prosopis glandulosa* (honey mesquite) which collectively dominate the vegetation community of the study site. The site is open to, but lightly used by domestic cattle.

### 2.2 Data collection

To quantify nurse-plant effects in our study system, we selected 30 pairs (15 for *L. tridentata* and 15 for *P. glandulosa*) of 30 × 30 cm plots in interspaces and under the canopy of neighboring shrubs; plots were on the northern aspect of each shrub. Shrub and interspace pairs were selected within a preselected polygon that avoided prior/ongoing research efforts and to avoid drainage channels, recent surface damage from livestock and wildlife, and to cover a range of shrub sizes while maintaining a minimum distance of ∼5 m between individuals (most shrub/interspace pairs were separated by ≥ 15 m). The same shrub/interspace pairs were subsequently used for soil biogeochemical sampling (see section 2.2.1). All plots were marked with stainless steel nails to enable accurate relocation and were repeatedly surveyed for emerging plants (approximately once per week for 4 to 6 weeks) during the monsoon seasons of 2020 and 2021. Emerging plants were identified to species whenever possible as part of a companion study on plant community assembly processes; all plants observed during this study were native to the ecosystem. None of the plants that emerged in 2020 flowered or survived (if perennial) beyond the fall season due to a prolonged drought. In contrast, an above average volume of monsoon rainfall in 2021 boosted performance and nearly 100% of emerging plants matured and flowered and/or survived through the year. We used paired tests, with one pair of plots in *L. tridentata* excluded because of persistent rodent digging, to verify that a greater number of plants occurred under shrub canopies than in interspaces each year (see section 2.3).

#### 2.2.1 Soil biogeochemistry

In late June of 2020, soil samples were removed with a one-piece probe sampler (1.9 cm diameter opening) to a depth of 10 cm for biogeochemical analysis. The samples were collected within a 1-hour period, placed into separate plastic bags, sealed into a cooler with ice, and shipped overnight to the USGS Southwest Biological Science Center in Moab, UT, for next-day processing. In the lab, samples were sieved through 2 mm mesh to remove rocks and large organic material and weighed into scintillation vials or tins within 3 hours of arrival for use in biogeochemical analyses. Soil organic matter was assessed by comparing pre- and post-combustion weights of samples heated to 550 °C for 4 hours in a muffle furnace (ThermoScientific Thermolyne, Waltham, MA, USA). Total (%) C and N were measured via sample combustion using a Vario MICRO Cube (Elementar Americas, Mt. Laurel, NJ). Measures of total soil-extractable organic C (TOC) and total dissolved N (TDN) were determined by adding 10 g of sample to 35 ml of 0.5 M K_2_SO_4_, mixing for 1-hour on an orbital shaker table, filtering on Whatman #1 filter paper, and analyzing the resulting solution with a Shimadzu TOC-VCPN with a TNM-1 + ASI-V total-N measuring unit. Soil-extractable ammonium (NH_4_ ^+^-N) and nitrate (NO_3_ ^-^-N) concentrations were measured by adding 10 g of sample to 35 ml of 2 M KCl, shaking for one hour and allowing solutions to sit overnight, filtering on Whatman #1 filter paper, and analyzing the resulting solution with a Smartchem 200 Discrete autoanalyzer (Milford, MA). Phosphorus extracted from soils was measured as PO_4_ ^3-^-P via bicarbonate extraction (Olsen & Sommers, 1982) and quantified via colorimetric reaction using an ascorbic acid molybdate analysis (Kuo, 1996) on an autoanalyzer (Westco Scientific Instruments, Waltham, MA). Microbial C and N concentrations were estimated with a chloroform cell lysis method by adding 1 ml of amylene-stabilized CHCl_3_ to 10 g soil in a 125 ml flask that was stoppered with neoprene and allowed to sit in the dark for 16 hours before being ventilated and extracted with 0.5 M K_2_SO_4_ (Beck et al., 1997; Brookes et al., 1985) and shaken for 1 hour. Microbial biomass C and N were calculated by subtracting the amounts of C and N extracted from nonfumigated soil from the amounts extracted from paired fumigated soil after analysis on a Shimadzu TOC-VCPN with the TNM-1 + ASI-V attachment (Shimadzu Corporation, Kyoto, Japan); no microbial biomass correction factors were applied (Weintraub et al., 2007).

#### 2.2.2 Solar radiation and soil temperature

Photosynthetically active radiation (PAR in mol m^-2^ s^-1^) reaching the ground-surface in unvegetated interspaces and beneath shrub canopies was measured between 5 August and 6 September 2020 using a set of 4 Apogee MQ-500 full-spectrum quantum sensors. During each deployment, one sensor was placed in an interspace and companion sensors were placed beneath three neighboring shrubs and collected PAR data at 1-minute intervals for 72 or more hours, after which, the sensors were relocated to a unique interspace and set of shrubs. This process was repeated until ground-level PAR had been recorded beneath the canopies of 12 *L. tridentata* shrubs and 18 *P. glandulosa* shrubs; interspace/shrub combinations were selected to cover a range of shrub sizes while minimizing the need to reposition a central data logger. We opted to measure fewer *L. tridentata* given its more uniform canopy architecture relative to the more variable canopies of *P. glandulosa* of this study system. Given the sampling design, variability in PAR reaching the surface of the unvegetated interspaces was overwhelmingly caused by changes in daily atmospheric conditions (i.e., cloud cover), and to a far lesser extent, variation from the changing angle of the Earth relative to the Sun. Thus, PAR values recorded beneath shrubs were considered in relation to values recorded in the neighboring interspace to account for atmosphere-caused and diurnal-caused variation in solar radiation. PAR values were recorded in µmol m^-2^ s^-1^ and values < 100 and > 2200 were excluded from the analysis of PAR to reduce bias caused by extreme low light times of the day and to remove extremely high values that include a substantial amount of radiation bounce-back from neighboring vegetation surfaces and soils. Values above 2200 were rare throughout the study while values below 100 were recorded daily from just before sunset to just post sunrise. Soil temperature in relation to PAR sensor placements was measured at 15-minute intervals with Hobo Pendent temperature sensors (Onset Computer Corporation, Bourne, MA, USA) placed horizontally into the soil between 5 and 7 cm deep. To avoid disturbing the soil directly above the sensors, a small soil pit was excavated with a hand trowel to allow the creation of a sensor-sized cavity at the desired depth into which each logger was pushed; the small pits were then refilled with the extracted soil to fully cover the sensors.

#### 2.2.3 Soil moisture

Soil moisture in our study site was measured at a depth between 5 and 7 cm as volumetric water content (VWC) using 18 Decagon ECHO-EC5 soil moisture sensors (Meter, Pullman, WA, USA) inserted horizontally into the soil. To avoid disturbing the soil directly above the sensors, a small soil pit was excavated with a hand trowel to allow the creation of a sensor-sized cavity at the desired depth into which each sensor was pushed; the small pits were then refilled with the extracted soil to fully cover the sensors. VWC values were recorded at 15-minute intervals between 30 June 2020 and 3 October 2021 in 6 unvegetated interspaces (4 open and 2 covered with shade cloth structures) and in soils under 6 shrubs each of *L. tridentata* and *P. glandulosa*, (4 intact and 2 with aboveground biomass trimmed away during the study; both species can resprout from root material and all shrubs, intact or trimmed, were verified to be alive throughout the entirety of our study). Shade cloth covers, used as mimics for shrub canopies, were verified to intercept a similar amount of radiation as shrubs and then employed in this study to assess the effect of shading on soil moisture without influences of shrub root systems that have been reported to ‘lift’ moisture hydraulically from deeper to near surface layers (Caldwell et al., 1998). The soil moisture sensors described above were installed in 2 locations of the larger study area that were roughly 25 m × 25 m in dimension in a western and eastern position that corresponded to “upslope” and “downslope,” respectively, along the 7 m cline in elevation of the study area. These locations were selected to aid in assessing spatial variation in soil moisture while dealing with logistical constraints imposed by wire lengths of the sensors and the need to position a central data logger for each set of sensors. All shrubs and interspaces selected in the two measurement areas were separated by a min of 4 and max of 25 m.

In addition to field measurements of soil moisture, we measured gravimetric water content experimentally in the lab. When the VWC sensors were removed from the study site on 3 October 2021, 3 soil samples, each roughly 1.5 L in volume, were collected from a depth of 5 to 7 cm in areas closely neighboring each sensor (surface material was pushed aside before collection). The soil samples were air dried and sifted through 4 mm mesh before being subsampled into plastic pots (9 × 9 cm opening and 23 cm depth) lined with nylon mesh to retain soil but allow for water drainage. Each pot was outfitted with 1 Decagon ECHO-EC5 soil moisture sensor, positioned vertically to cover soil depths ranging from 3 to 8 cm. Two laboratory trials were completed—1 trail each for soils collected from the upslope and downslope measurement locations of the field site—between January and April 2022. Each trial consisted of 18 pots, 6 each for interspace, *L. tridentata*, and *P. glandulosa* soils. Distilled H_2_O was added to the surface of each pot in ∼ 50 ml aliquots, with subsequent aliquots added once surface ponding had disappeared, until water began to drip from drain holes on the bottom of the pot. After water began draining from the bottom of each pot, they were allowed to sit for 1 hour and were then weighed; this 1-hour value was later used to calculate the maximum water content (i.e., content at lab saturation) for each mesocosm. Pots where then weighed every 6 to 8 hours for 3 to 4 days and then weighed every 12 to 24 hours for the remainder of each trial to allow for calculation of dry down rates and to generate timeseries of VWC values and pot weights for calibrating the Decagon ECHO-EC5 sensors (below). At the conclusion of each dry-down trial, VWC sensors were removed and mesocosms were placed into an oven at 70 °C for ≥ 5 days before being weighed; dry soil weight for each pot was then calculated by subtracting the weight of the empty plastic pot from the post-oven dried value. Water weight in each mesocosm during the recorded values of the dry-down trials was subsequently calculated as: H_2_O (g) = mescocosm weight in time series (g) – [dry soil (g) + plastic pot (g) + VWC sensor (g)].

Values of water weight and sensor-derived VWC were used to calibrate the ECHO-EC5 sensor data from the field setting; spatial variation was accounted for by georeferencing the soil samples of lab mesocosms to the ECHO-EC5 that closely neighbored the collection location in the field site. Given differences in max saturation and dry-down rates, separate mixed-effect models with random intercept terms for soil sensor ID to account for repeated measures, was created for soils of interspaces, *L. tridentata*, and *P. glandulosa*. All models included the laboratory derived ‘true VWC’ based on mesocosm wet – dry weight as the dependent variable and sensor-derived VWC and the field collection location as fixed-effects. The models were strong fits to the data with marginal R^2^ ≥ 0.93 for all three. Parameter values of each model (Table S#) where then used to hindcast and correct field-based VWC values of ECHO-EC5 sensors in the R statistical environment (R Core Team, 2022).

### 2.3 Data analyses

All data processing and analyses were completed in the R environment using RStudio version 2022.07.2 and R version 4.2.1 (R Core Team, 2022). Scripts are available upon request. Difference in seedling/plant occurrence in paired shrub/interspace plots were tested with paired mixed models, where the random intercept term was pair ID and fixed effects included cover (i.e., interspace or shrub) and year (i.e., 2020 or 2021), using the ‘lmer’ function of the ‘lme4’ package (Bates et al., 2015).

#### 2.3.1 Soil biogeochemistry data analyses

We used Moran’s I to test for but did not find evidence of significant spatial autocorrelation across the study plots for any of the biogeochemical measures considered (P > 0.05 for all). Thus, we dismissed geographically weighted models for analyses. We used PERMANOVA to compare the multivariate biogeochemical metrics of interspace soils located near *L. tridentata* and *P. glandulosa* shrubs, values from under the two species, and values among interspaces and shrubs for both species. We used the ‘effectsize’ package to calculate paired Mahalanobis’ D and Cohen’s *d* values to compare effect sizes among the species (Ben-Shachar et al., 2020). We also used PERMANOVA to compare concentrations of the individual biogeochemical variables of interspace and shrub soils followed by pairwise, post-hoc comparisons when the omnibus model had P ≤ 0.05. All PERMANOVA modes were completed using the ‘adonis2’ function of the ‘vegan’ package (Oksanen et al., 2013) and post-hoc tests were completed with the ‘pairwise.perm.t.test’ of the ‘RVAideMemoire’ package (Hervé, 2020). The influence of shrub size, measured as areal cover in m^2^, was considered in permutational linear models for each of the 12 biogeochemical variables for both species using the ‘lmp’ function of the ‘lmPerm’ package (Wheeler & Torchiano, 2016).

#### 2.3.2 Solar radiation and soil temperature data analysis

We considered the influence of shrub species, canopy cover, and incoming solar radiation at the time of measurement on reduction in levels of PAR reaching the ground surface beneath shrubs and on soil temperature using linear mixed-effects models with a random intercept term for sensor ID to account for repeated measures over time. The models for PAR and soil temperature were modeled with a Gaussian distribution and specified using the ‘lmer’ function of the ‘lme4’ package (Bates et al., 2015). In total, we considered 3 models for PAR (see Table S2 for model specification): (1) a comparison of PAR reaching the ground surface in interspaces and beneath shrubs, (2) an assessment of the influence of shrub species and canopy cover on the relative change in PAR beneath shrubs relative to interspaces, and (3) the response of PAR recorded under shrubs in response to incoming PAR and shrub species/canopy cover. For the second PAR model above, relative change in PAR was calculated as the standardized difference of PAR reaching sensors placed beneath shrubs and those in unvegetated interspaces (i.e., RCI_PAR = [PAR_shrub_ – PAR_interspace_] / PAR_interspace_). We compared 4 candidate models each for soil temperature (interspaces and beneath shrubs) and soil temperature under shrubs only (see Table S2 for model specification). Visualization of soil temperature as a function of incoming PAR revealed non-linear relationships in the data, leading us to fit 2 of the 4 candidate models for the full-set and sub-set of soil temperature data (Table S#) with flexible splines using the ‘splines’ package in base R. The ‘best-fit’ model from each set of 4 candidates was determined using AICc values calculated with the ‘performance’ package (Lüdecke et al., 2021). The data were limited to times when unobstructed PAR was > 100 and ≤ 2200 mol m^-2^ s^-1^ to avoid biasing the models by removing measures from times when the sun was below or very near to the horizon and to reduce times when radiation bounce-back from neighboring vegetation was very high. We used the ‘rstatix’ package (Kassambara, 2023) to calculate Cohen’s *d* values and 95% confidence intervals (using the adjusted bootstrap percentile or ‘bca’ method) of effect size for reduction of PAR and soil temperatures for each shrub species relative to interspaces at an hourly scale between 0600 and 2000 hours—the window of time corresponding to the hours between sunrise and sunset during this study.

#### 2.3.3 Soil moisture data analysis

In the laboratory dry-down experiment, water volume at maximum saturation of soils collected from interspaces versus beneath the shrub species were compared via two-way permutational ANOVA with soil provenance (interspace, *L. tridentata*, and *P. glandulosa*) and collection location (eastern vs. western collection location, see section 2.2.3) as the independent terms using the ‘lmp’ function of the ‘lmPerm’ package (Wheeler & Torchiano, 2016). The proportion of max saturation water volume and VWC over time of soils collected from interspaces versus beneath the shrub species in the laboratory dry-down experiment was analyzed using linear mixed-effect models with a Gaussian distribution and specified using the ‘lmer’ function of the ‘lme4’ package (Bates et al., 2015). Both models had soil provenance, collection location, and time since max saturation occurred as fixed effects and a random intercept term for mesocosm ID to account for repeated weight and VWC values.

Prior to analysis of the field site VWC data, values < 0.0032 (lowest 1 % of all values) were considered to be outliers and were removed from the time series of all sensors; given their design, ECHO-EC5 sensors still record positive values even when VWC = 0 in the soil matrix. VWC values were then square root transformed to approximate a Gaussian distribution and analyzed using linear mixed-effect models specified using the ‘lmer’ function of the ‘lme4’ package (Bates et al., 2015). The model included cover type (intact shrub, shade cloth, no cover, and trimmed shrub), shrub species (either over the sensor or neighboring the interspace location), and sensor position in the study site (eastern vs. western location); see section 2.2.3 for description of VWC sensor placement. The VWC time series for each sensor were also analyzed for water ‘pulses’ which represented an increase in VWC (caused by precipitation) above a threshold; pulses were identified using the method described below which allows for pulses to persist for multiple days in the time series. After identifying and quantifying the duration of VWC pulses, we computed two-way permutational ANOVAs—using the ‘lmp’ function of the ‘lmPerm’ package (Wheeler & Torchiano, 2016) for the mean number of pulses per day (pulses recorded / days in the time series) and the proportion of time that VWC was above the threshold for a pulse event in each time series. Each model had sensor position in the study site (eastern vs. western location) and sensor location—i.e., location within the 6 primary cover groups that included interspace open and shaded, *L. tridentata* intact and trimmed, and *P. glandulosa* intact and trimmed.

To identify ‘pulses’ in VWC recorded by each sensor, a pulse detection algorithm based on z-scores was applied to each sensors’ time series (see Brakel, 2014). We applied the algorithm to a smoothing spline fit to the time series data of each sensor with the ‘smooth.spline’ function of the ‘stats’ package in base R. Fitting a spline removed the influence of diurnal variation in VWC caused by temperature sensitivity of the ECHO-EC5 while applying the algorithm to each sensor’s time series allowed local baselines and variation in the data determined whether VWC was above the threshold for being considered a pulse. This approach allows for a robust comparison of relative soil VWC variation among sensors by subjecting each time series to similar criteria for detecting pulses while using local soil VWC values to set the threshold for detecting a pulse—a method that subjects wetter and drier locations to a similar test. In brief, the algorithm constructs a separate moving mean and deviation, such that signals do not greatly alter the threshold for identifying a pulse, allowing future signals to be identified with approximately the same accuracy, regardless of the number and size of previous signals. The algorithm utilizes 3 inputs: (1) the time lag of the moving window, (2) a z-score standard deviation threshold for determining the start and end of pulse events, and (3) the influence (between 0 and 1) of new signals on the mean and standard deviation. The time lag was set to 168 hours (1 week), influence of new signals was set to 0—which assumes time series stationarity—so that each pulse was evaluated on the same criteria, and the threshold was set to 2.5—a value that corresponds to the top 1.25 % of VWC values in the time series of each sensor. VWC sensors were installed into the study site approximately 2 days prior to a large rain event that initiated the seasonal monsoon period of this ecosystem. To provide a baseline of VWC values that reflected background conditions for each VWC sensor location, we appended 1000 hours of simulated VWC values drawn from a theoretical Gaussian distribution to the start of each time series for training the algorithm on relatively dry soils. The values were generated using the ‘rnorm’ function of the ‘stats’ package in base R using the mean and standard deviation of values ≤ to the 75^th^ quartile of the original data time series of each VWC sensor—an approach that removes the top 25% of VWC values to train the algorithm on realistic values for drier periods of this system.

## 3. Results

Significantly more seedlings/plants were observed under shrub canopies than in interspaces of the paired plots (P < 0.0001, Figure S1b) and significantly more plants were observed in the above average rainfall year of 2021 compared to the below average year of 2020 (P < 0.0001, Figure S1b).

### 3.1 Soil biogeochemistry

The multivariate soil biogeochemical pools beneath *L. tridentata* and *P. glandulosa* did not significantly differ (Figure S3, P = 0.084, F = 2.33, and d.f. = 1, 28), nor did the pools of the unvegetated interspaces that were paired with *L. tridentata* versus *P. glandulosa* shrubs (P = 0.686, F = 0.491, and d.f. = 1, 28; Figure S3). The biogeochemical pools of soils from interspaces versus beneath shrubs differed significantly for both species (P = 0.001, F = 17.41, d.f. = 1, 28 and P = 0.001, F = 18.30, d.f. = 1, 28 for *L. tridentata* and *P. glandulosa*, respectively; Figure 2). Effect size differences among the biogeochemical pools of interspaces and beneath shrubs were similar for both species, with Mahalanobis’ D = 2.82 (95% CI = 0.79, 3.13) and 2.88 (95% CI = 0.85, 3.19) for *L. tridentata* and *P. glandulosa*, respectively.

**Figure 2.**
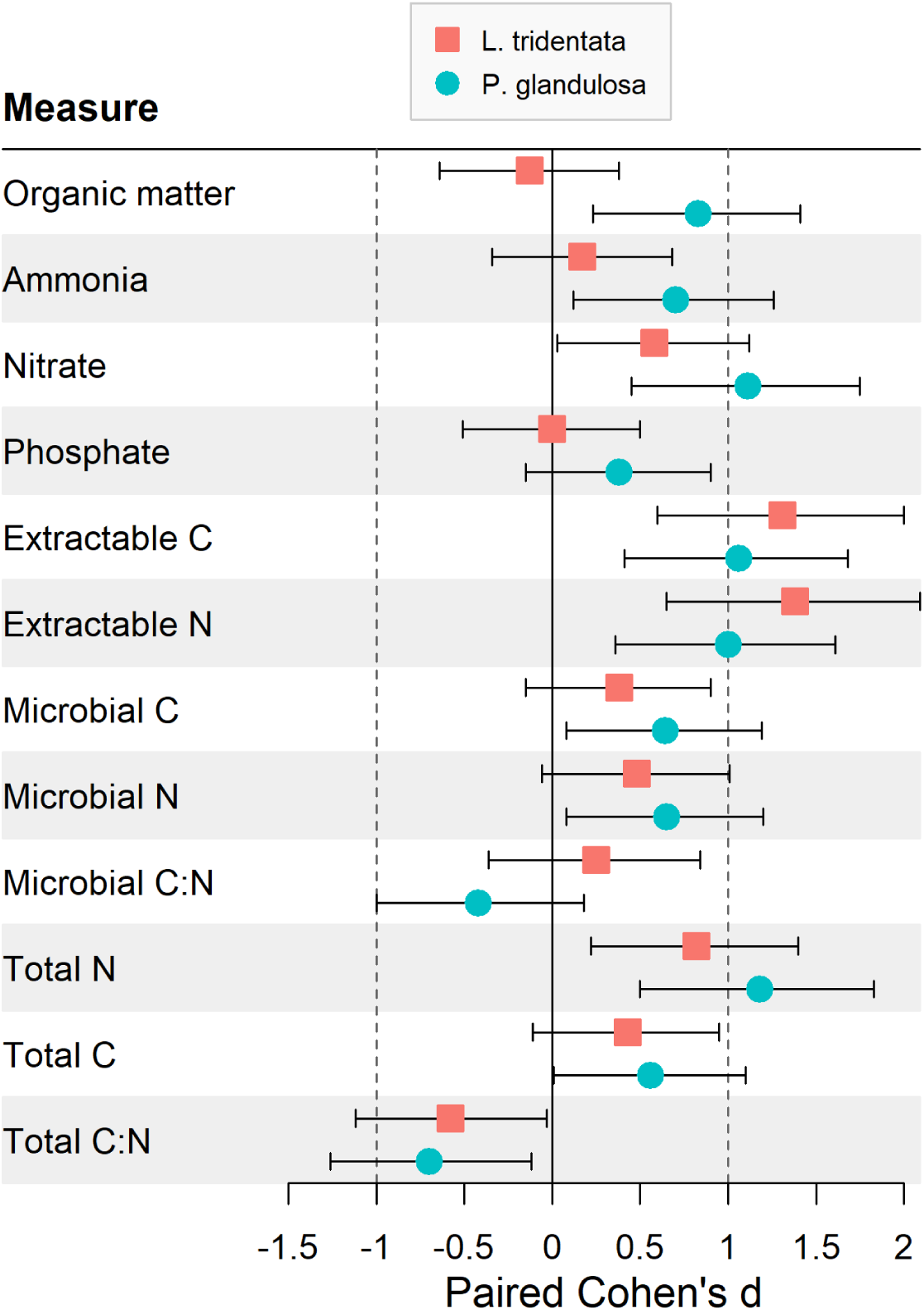
Paired Cohen’s *d* values (and 95% confidence intervals) quantifying the “shrub effect” of *L. tridentata* (teal circles) or *P. glandulosa* (red squares) on individual biogeochemical variables in soils. Values indicate effect size differences between values in soils sampled beneath shrubs compared to soils from neighboring, unvegetated interspaces paired with each shrub. Positive values indicate a higher value was measured in soils beneath shrubs relative to the interspace, negative values indicate the opposite.

Of the 12 biogeochemical variables assessed, six were significantly influenced by shrub species (organic matter, Total N, extractable N, NH_4+_, NO_3-_, and PO_4+_) with higher values found in soils under *P. glandulosa* than under *L. tridentata* (Figure 2 and Figure S4). Only one variable—extractable N— significantly differed among interspaces from *L. tridentata* versus *P. glandulosa* sampling blocks (P < 0.05, Figure S4). A paired Cohen’s *d* with 95% confidence intervals was used to measure the effect size of shrubs—i.e., the “shrub effect”—relative to interspace soils in each shrub-interspace pair for both species. Eight biogeochemical variables—including organic matter and all measures of soil N—were found in significantly greater concentrations in soils under *P. glandulosa* versus paired interspaces while the total C:N ratio was significantly lower under the shrub (Figure 3). Four variables were found in significantly greater concentrations in soils under *L. tridentata* versus paired interspaces while only the total C:N ratio was significantly lower under the shrub (Figure 3). For both shrub species, forms of soil N comprised the majority of the variables that were significantly greater in soils collected from under the shrubs relative to soils collected from the neighboring, paired interspace. Shrub size in *P. glandulosa*, measured as canopy volume, had a significant positive effect on the concentration of four of the 12 biogeochemical variables (NO_3_ ^-^, microbial biomass C and N, and total N) while the size of *L. tridentata* did not have an effect on any biogeochemical variables (Figure S6).

**Figure 3.**
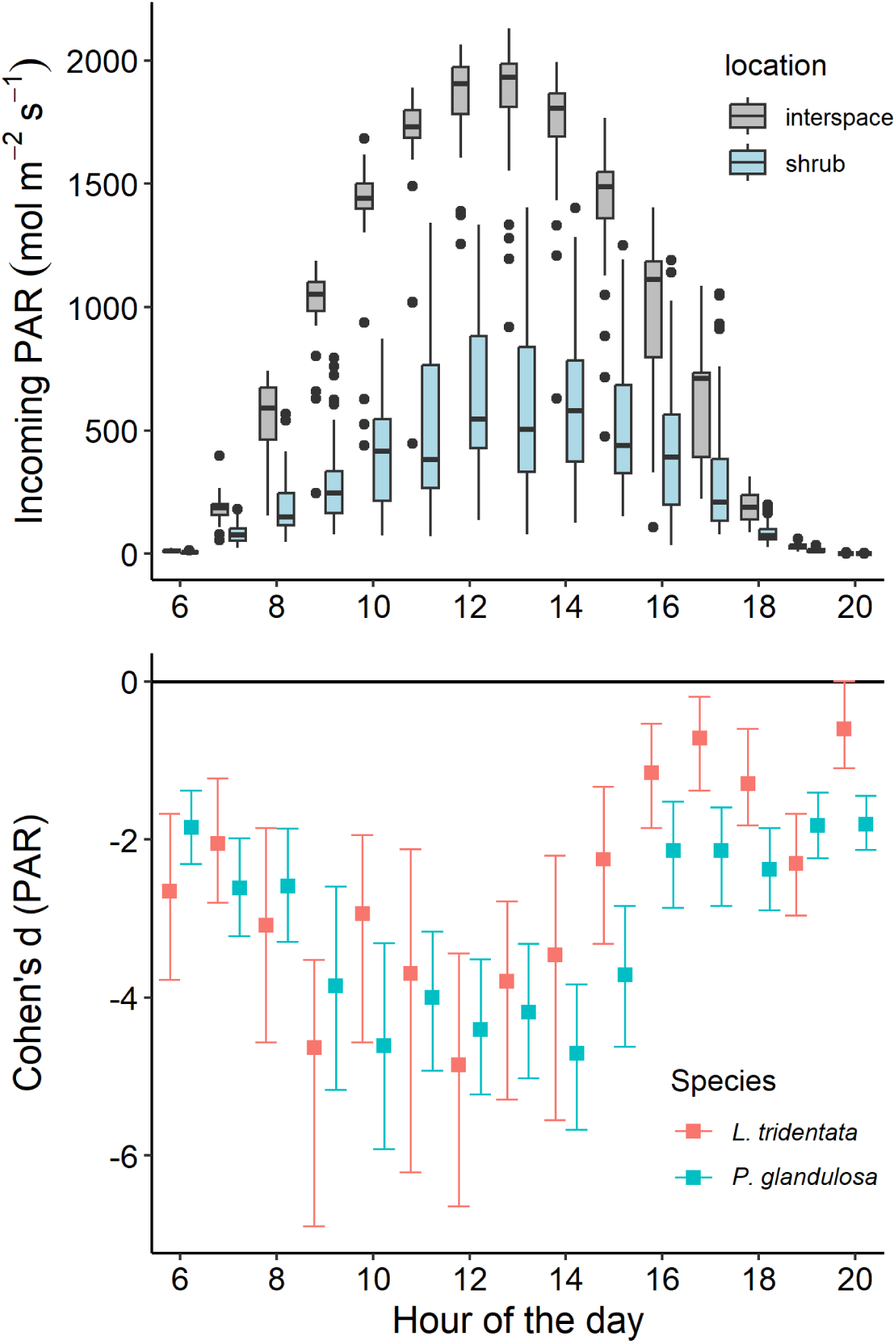
Comparisons of incoming photosynthetically active radiation (PAR) measured in unvegetated interspaces and under neighboring shrubs of *L. tridentata* and *P. glandulosa* (top panel) and Cohen’s *d* effect size values, with upper and lower 95% confidence intervals (bottom panel), illustrating the effect of shrub cover on PAR reaching the ground surface under shrub canopies versus unvegetated interspaces. When included in a mixed model analysis, shrub species was not considered to be a significant influence (P > 0.05) on the reduction in PAR caused by aboveground biomass; mean hourly effect sizes of the two species differed only for two out of 24 hours—i.e., 17:00 (5:00 PM) and 20:00 (8:00 PM).

### 3.2 Solar radiation and soil temperature

Incoming solar radiation—measured as photosynthetically active radiation (PAR, mol m^-2^ s^-1^)— reaching the ground level beneath shrub canopies differed significantly from levels of PAR reaching the surface of unvegetated interspaces (P < 0.0001, Figure 3). The relative change in PAR, calculated as the standardized difference of PAR reaching pairs of sensors placed beneath shrubs and those in unvegetated interspaces (i.e., RCI = [PAR – PAR] / PAR), had a mean of −0.63 and ranged from −0.97 to 0.00 in value across the study period (Figure S#). Levels of PAR measured beneath shrubs was significantly and positively related to levels of incoming PAR at the time of measurement (P < 0.0001, conditional R^2^ = 0.38, marginal R^2^ = 0.31; Figure S#). The reduction in PAR caused by shrubs was related to canopy cover of shrubs (P = 0.043, Figure S#) but was not significantly different among the two species (P = 0.713). While the two factors had opposing influences, the effect of incoming PAR to the system at the time of measurement had an effect 4.19 times greater than canopy cover in shaping the amount of PAR reaching the ground surface beneath shrubs—a result that highlights the importance of simple shrub canopy presence in addition to the moderating effect of total canopy cover. The relationship among shrub canopy area (a correlate of overall shrub size) and canopy cover (proportional cover of the sky in upward facing photos) was slightly negative but not significant in *L. tridentata* (R^2^ = 0.14, P = 0.228) and significantly positive but weakly related in *P. glandulosa* (R^2^ = 0.30, P = 0.018) (Figure S#).

Soil temperature, recorded at a depth between 6 and 8 cm, was significantly and positively related to incoming solar radiation measured as PAR (P < 0.0001, Figure 4). Location of the soil temperature sensor —in terms of open interspace versus beneath shrub canopies—was not a significant effect (P = 0.25) on temperature but the interaction of location with incoming PAR was significant (P < 0.0001). The best-fit model included a flexible spline fit among PAR and temperature to account for the reduction in PAR reaching the ground surface under shrub canopies (conditional R^2^ = 0.53, marginal R^2^ = 0.47; Figure 4, Table S#). Sub-setting the data to include only soil temperatures beneath shrub canopies revealed no significant effect of species or canopy cover on soil temperature (P = 0.66 and P = 0.44, respectively) but, again, a significant effect of incoming PAR and an interaction among species and PAR on soil temperature (P < 0.0001 for both, conditional R^2^ = 0.49, marginal R^2^ = 0.41). The reduction in radiation and the related reduction in the effect size of soil temperature beneath shrub canopies relative to interspaces was qualitatively similar throughout the day among the two species but with a greater number of hours on average throughout the day (i.e., 11 vs. 8) being significantly cooler in shrub versus interspace soil under *P. glandulosa* than *L. tridentata* (Figure 5).

**Figure 4.**
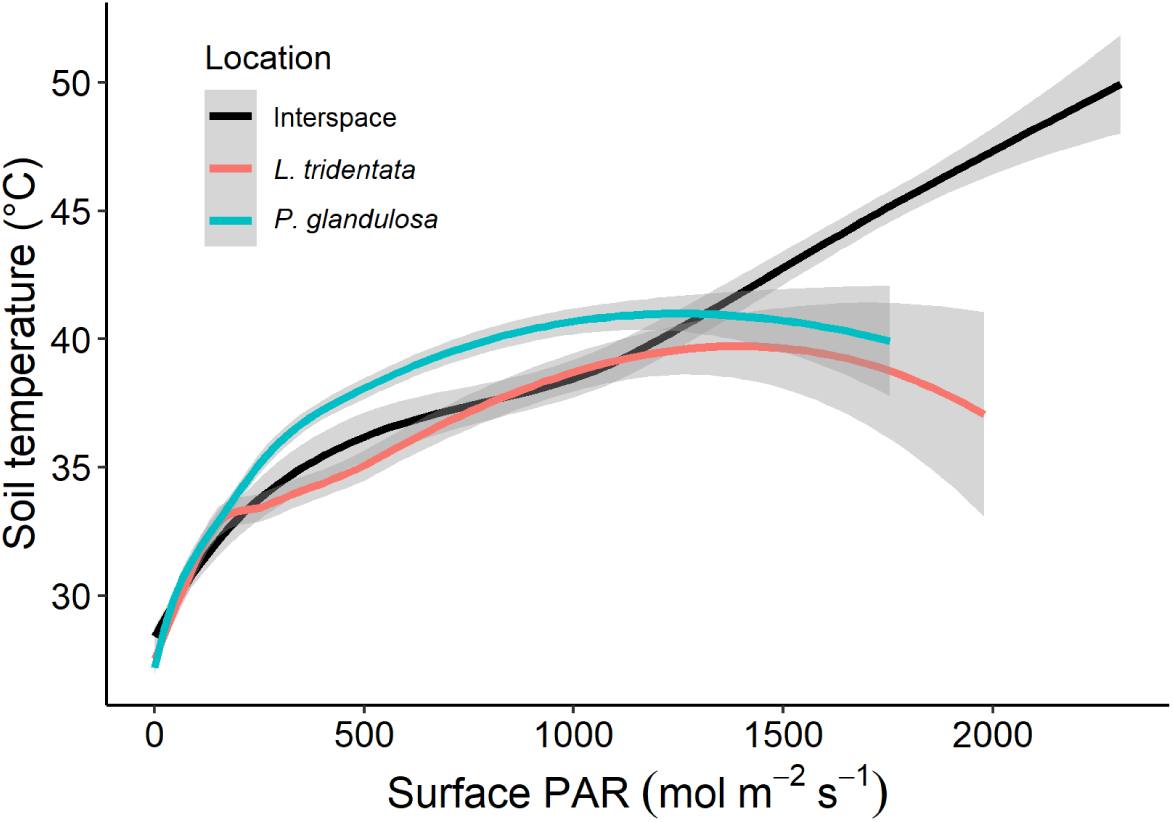
Soil temperature, between 6 and 8 cm depth, as a function of incoming solar radiation within the photosynthetically active radiation (PAR) spectrum in soils of unvegetated interspaces or beneath the canopy of *L. tridentata* or *P. glandulosa*.

**Figure 5.**
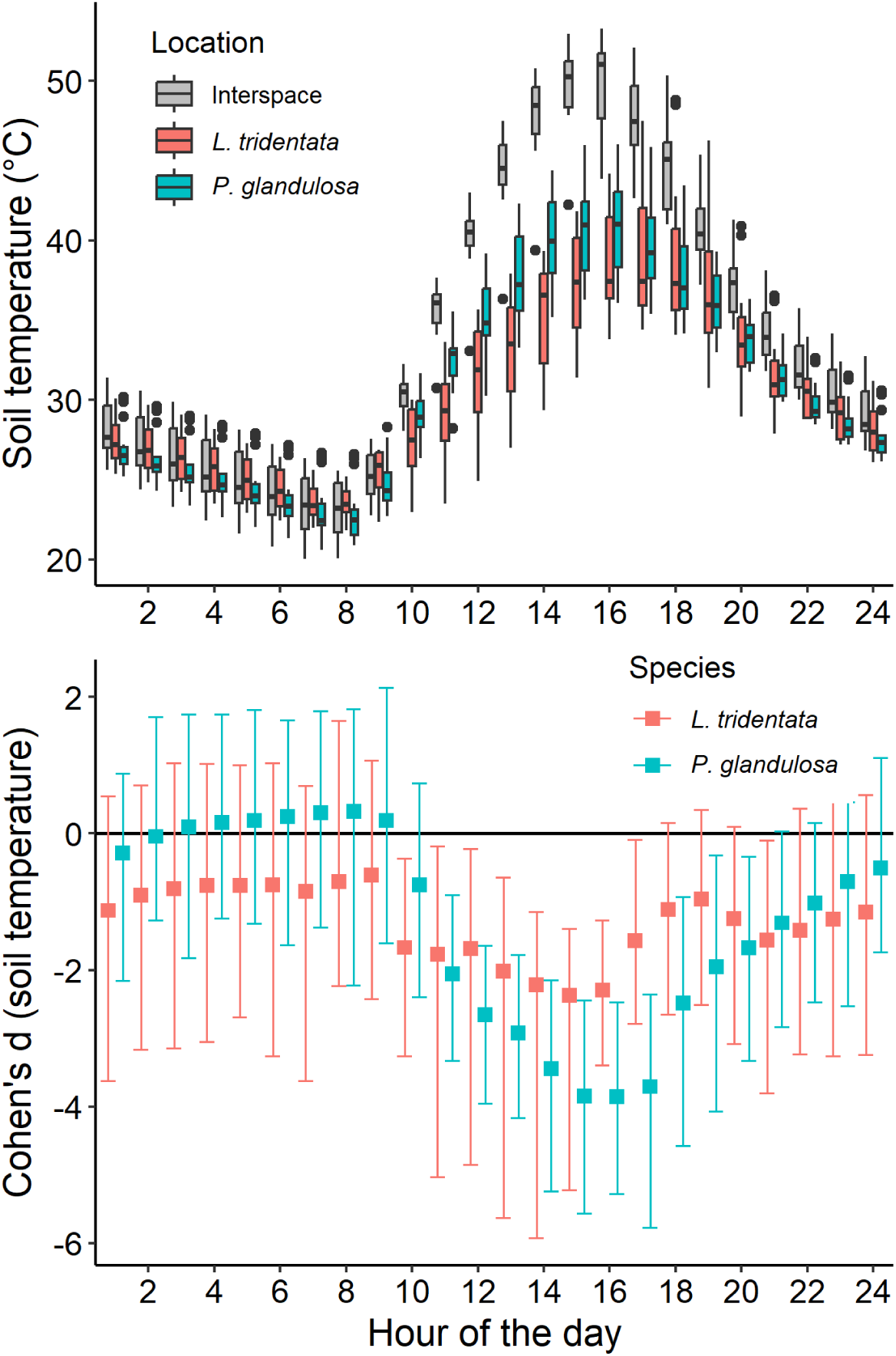
Mean hourly soil temperature, between 6 and 8 cm depth, throughout the study period (top panel) within unvegetated interspaces or beneath the canopy of *L. tridentata* or *P. glandulosa*. (bottom panel) Cohen’s *d* values (and 95% confidence intervals) quantifying the “shrub effect” of *L. tridentata* or *P. glandulosa* on soil temperature; values indicate effect size differences between measures in soils beneath shrubs compared to soils from neighboring, unvegetated interspaces with negative values indicating a lower temperature beneath shrubs relative to the interspace, positive values indicate the opposite.

### 3.3 Soil moisture

At max saturation, soils collected from beneath *P. glandulosa* held a significantly greater volume of water (P < 0.0001) and had a larger VWC value (P < 0.0001) relative to soils sourced from *L. tridentata* and unvegetated interspaces (Fig. 6). Over time, following max saturation, soil VWC was significantly affected by the amount of time that had passed (P < 0.0001, Figure 6, Table S6) and also by soil provenance both in terms of its source location in relation to interspaces and shrubs (P < 0.0001) and whether it had been collected from the eastern versus western portion of the study site (P = 0.0002) (conditional R^2^ = 0.93, marginal R^2^ = 0.89). Similar to VWC, the proportion of water volume at max saturation remaining over time had a significant negative relationship to time since saturation (P < 0.0001) and was significantly greater in the shrub soils compared to interspace (P = 0.0090) but was not affected by soil collection location in the study site (conditional R^2^ = 0.94, marginal R^2^ = 0.88). Collectively, these values indicate that while soils under *P. glandulosa* shrubs can hold a greater volume of water than soils from beneath *L. tridentata* and open interspaces, the rate at which water is lost per unit of drying time after saturation remains similar for the two shrub species under laboratory conditions.

**Figure 6.**
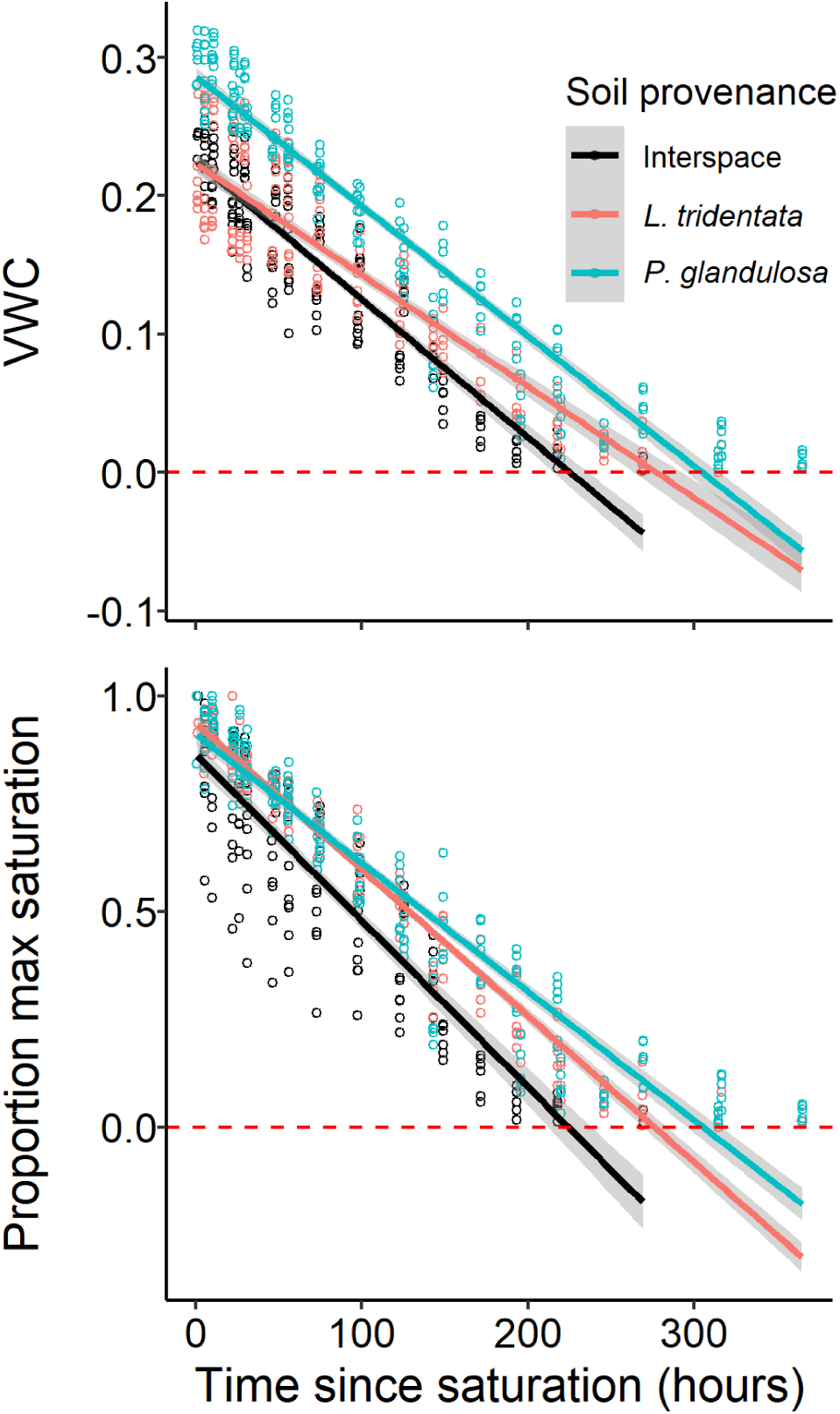
Soil moisture values—water remaining in grams (top panel) and proportion of maximum saturation water volume (bottom panel)—as a function of time since soil saturation in laboratory dry-down trials. VWC is volumetric water content.

In the field setting, VWC was significantly influenced by the location of sensors with regard to placement in interspace versus shrub soil, as well as location (upslope vs. downslope site) within the study site (R^2^c = 0.70, R^2^m = 0.59, Figure 7). Of the factors considered, location in relation to shrubs and interspaces was the most important factor, accounting for 89.0 % of the variance explained by the fixed effects in the model while local cover (i.e., open/trimmed vs. shaded/beneath shrub canopy) did not have a significant effect on VWC (Table S#). A *post-hoc* comparison indicated significant differences in the VWC of soils for all comparisons among interspace, *L. tridentata*, and *P. glandulosa* (P < 0.01 for all, Figure 7).

**Figure 7.**
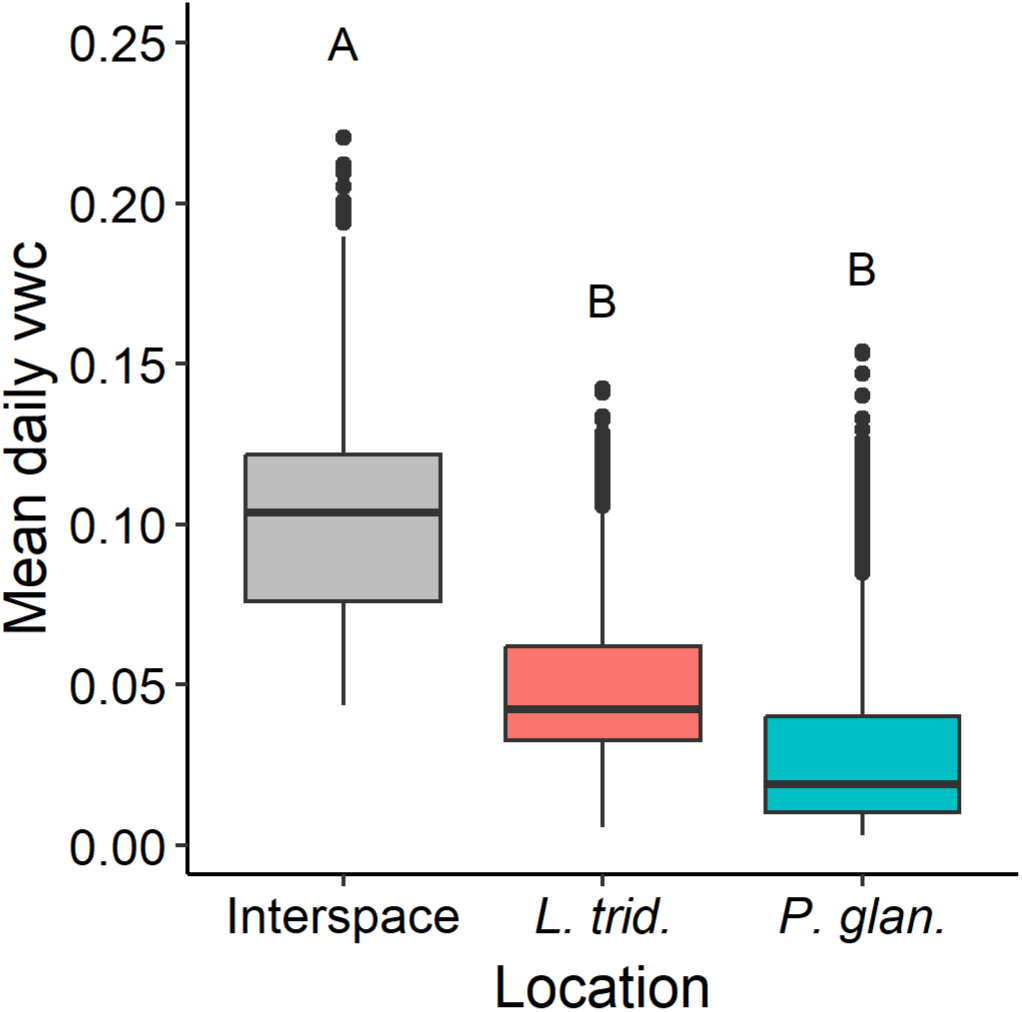
Soil volumetric water content (VWC) of the field site recorded by sensors located in the near-surface soils of unvegetated interspaces or beneath the canopy of *L. tridentata* or *P. glandulosa*. Cover treatments, including the addition of shade cloth to interspaces and the removal of aboveground biomass from shrubs did not have a significant effect on VWC.

Volumetric water content varied throughout the study in response to the timing and size of rainfall events (Figure S1). A comparison of the number of times across the hourly time series that VWC exceeded the threshold to be considered a “pulse event” indicated no significant differences among the soil locations or cover types (P = 0.41) but did reveal that VWC pulses were more common, on average, where canopy cover or shade cloth were absent (i.e., open interspaces and trimmed shrubs; Figure 8). When assessing the proportion of the time that VWC crossed the threshold and remained above it as part of variable duration pulse events, interspaces experienced a significantly greater proportion of time in pulse conditions compared to *L. tridentata*, and *P. glandulosa* (P < 0.05) while cover (i.e., open vs. covered by shrub biomass or shade cloth) did not have a significant effect (P > 0.05, Figure 8). The proportion of time above the pulse threshold among the locations mirrored values of mean daily VWC (Figure 7) with unvegetated interspaces exceeding the threshold for a larger proportion of time followed by *L. tridentata* and then *P. glandulosa*.

**Figure 8.**
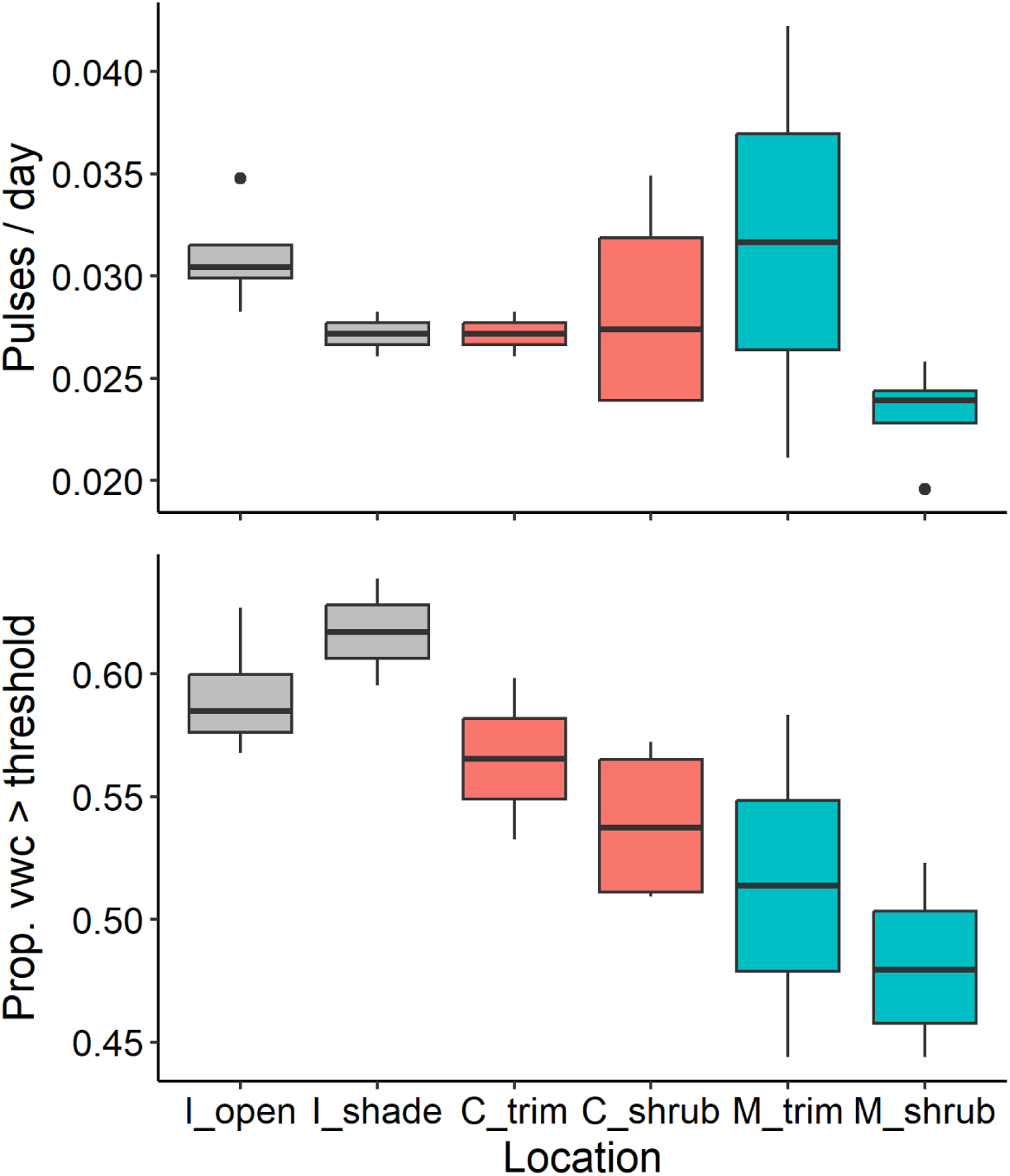
Using the time series logged by volumetric water content (VWC) sensors, the number of pulse events per hours in the time series (top panel) and proportion of the time series when VWC exceeded the threshold to be considered a pulse (bottom panel) for unvegetated interspaces (“I”) that were either left open or covered by a shade cloth roughly 30 cm above the ground surface that reduced radiation to a degree similar to shrubs (“shade”), or beneath the canopy of *L. tridentata* (creosote, “C”) or *P. glandulosa* (mesquite, “M”) that were either left intact (“shrub”) or trimmed at ground level to remove aboveground biomass (“trim”).

## 4. Discussion

Perennial plants, particularly shrubs, are often said to play the role of ‘nurse-plants’ in dryland ecosystems where they create ‘nurseries’ for neighboring/understory protégé plants that would otherwise be absent or occur at lower frequency/density without the environmental modifications caused by larger plants. Numerous mechanisms, operating individually or collectively, can create a nurse-plant effect (Filazzola & Lortie, 2014). A review of 158 reports from arid/semiarid systems indicated that the strongest likely drivers of positive nurse-protégé interactions were, in order of importance, 1) seed trapping by the nurse plant, 2) enhanced soil nutrient availability, 3) protection from herbivores/tramplers, and 4) increased water availability (Flores & Jurado, 2003). In this study, we found that two dominant shrub species of the Southwestern U.S., *P. glandulosa* and *L. tridentata*, harbored a greater number of protégé plants relative to neighboring interspace soils in both exceptionally dry and wet years. We simultaneously quantified several potential drivers of the observed nurse-plant effect including soil nutrient and water availability, and soil temperature, with a focus on comparing the effect each shrub species had on these drivers.

We found support for our first and second hypotheses that the nurse-plant effect in this system resulted, at least in part, from a combination of increased soil nutrient availability and a reduction in solar radiation and soil temperature beneath shrub canopies relative to unvegetated interspaces (Figure 9). Despite the reduction in solar radiation and soil temperatures beneath shrubs, we did not find support for our third hypothesis that soil available moisture would increase beneath shrub canopies. Instead, VWC sensors recorded greater soil moisture in unvegetated interspaces. This outcome contradicts reports from numerous dryland systems (e.g., Hong-Fei et al., 2010; Kidron & Gutschick, 2013; Li et al., 2010) and appeared to be driven by local, small-scale topography and surface flow dynamics of water during precipitation events. Importantly, shrub size was significantly related to the level of reduction in radiation and soil temperature—and size in *P. glandulosa* had a positive effect on some soil biogeochemical pools (Figure 9)—indicating that the strength of the nurse-plant effect provided by these foundational shrubs not only varies among individuals of different sizes, but also the importance of some mechanisms can increase over time as individuals grow larger. The relative importance of any mechanism driving a nurse-plant effect may vary at both short (diurnal/seasonal) and long (interannual/decadal) temporal scales. The dynamic nature of drivers of the nurse plant effect creates a need to quantify specific mechanisms and their combined influence over wider spatial and temporal scales if we are to advance our understanding of how perennial plants shape ecological processes and communities in drylands at landscape scales.

**Figure 9.**
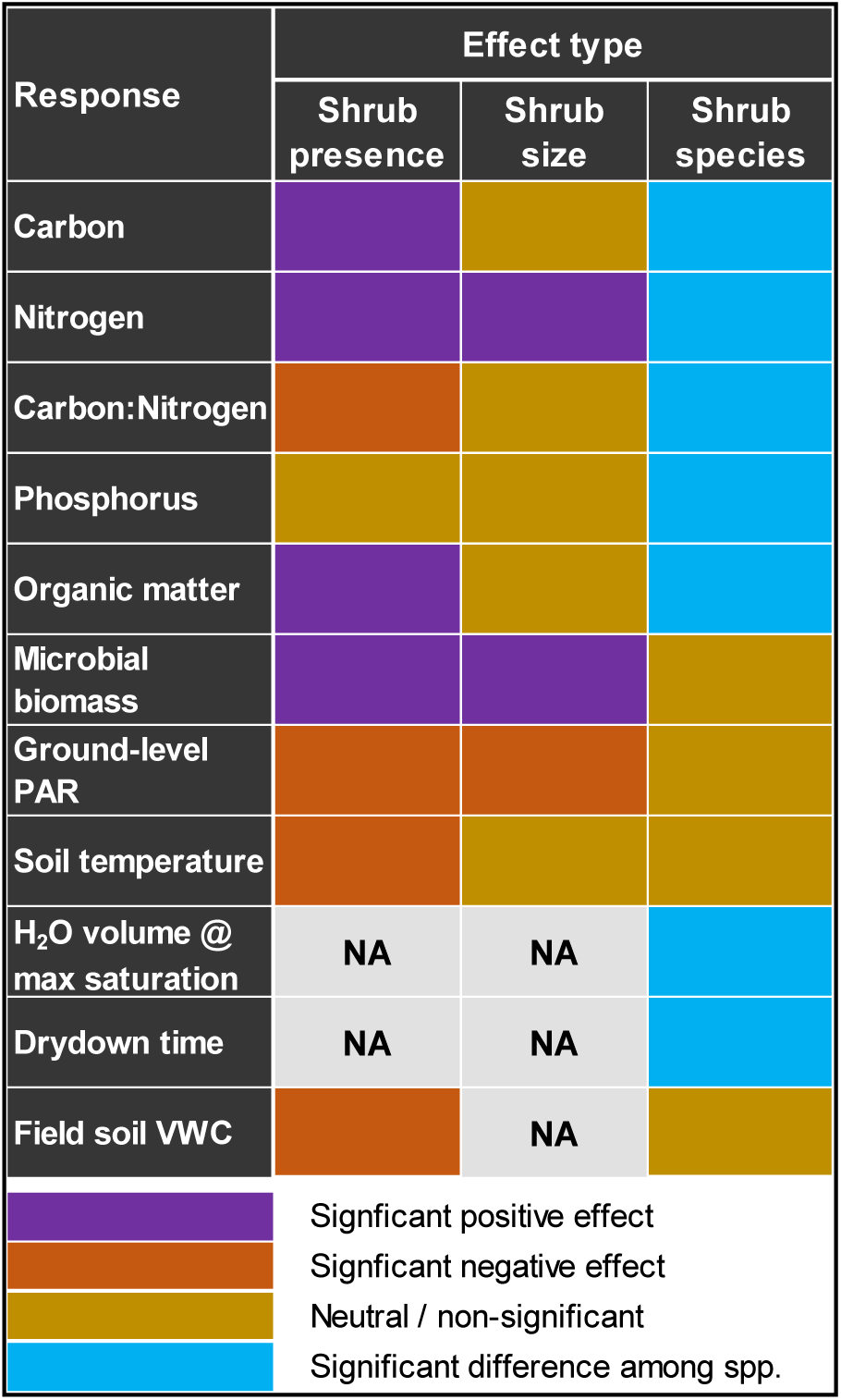
Summary of shrub effects on soil biogeochemical pools, solar radiation (PAR), and soil temperature and moisture values. Shrub effects are parsed here as presence/absence effects (i.e., an effect of shrub presence relative to unvegetated interspace values), shrub canopy size or area effects, and shrub species effects (i.e., significant difference in values between *L. tridentata* and *P. glandulosa*). Cells marked as having significant effects in the “shrub presence” and “shrub size” columns indicate at least one of the shrub species had values that significantly differed from the unvegetated interspace. VWC is volumetric water content. Grey boxes indicate potential relationships that were not assessed in this study.

### 4.1 Soil biogeochemistry

Plants can modify soil conditions in drylands by increasing carbon and nutrient concentrations and retention via root exudates, litter inputs, and microclimatic modifications (Schlesinger et al., 1996). As grass and forb cover declines in response to global change pressures affecting drylands, shrub cover has increased and interspace areas with sparse to zero plant cover have expanded in size and extent. This process has produced spatially discrete patches beneath/around shrubs where soil nutrients are found in far greater density relative to adjacent interspaces (Peterjohn & Schlesinger, 1990; Schlesinger et al., 1996). We found both species enhanced soil N and C pools but that *P. glandulosa* had broader and larger effects on soil biogeochemistry than *L. tridentata*, particularly with regard to the concentrations of total and inorganic N and organic matter (Figures 2, 9, S3, S4, and S5). Similar patterns for soil N and C pools have been previously reported for the Chihuahuan Desert in addition to other dryland ecosystems (Gao et al., 2022; Schlesinger & Pilmanis, 1998). Soil P is often a limiting or co-limiting element of plant growth in drylands (Drenovsky & Richards, 2004; He et al., 2014; James et al., 2005) and availability depends on mineral source from parent material or dust and/or the recycling of organic matter (Porder & Ramachandran, 2013). We found soil concentrations phosphorus were greater under *P. glandulosa* than under *L. tridentata* but that they were not significantly elevated relative to neighboring interspaces for either species. This result contrasts with prior work in nearby areas of our study system where soil P was found to be strongly linked to the extent of shrub canopies in *L. tridentata* (Schlesinger & Pilmanis, 1998).

The distribution of soil nutrients, including P, can be shaped by a variety of abiotic and biotic factors that can correlate with but also operate independently of shrub cover. For example, animal burrowing, erosion and sedimentation, water flow along the ground surface or plant stems and branches, and surface biotic community type and composition have all been demonstrated to affect soil nutrient concentration and distribution (Crain et al., 2018; Porder & Ramachandran, 2013; Schlesinger & Pilmanis, 1998). In our study system, cover by biological soil crusts (biocrusts)—communities of non-vascular organisms that aggregate mineral soils into a coherent surface layer (Weber et al., 2022)—has been linked to a reduction in soil P concentrations relative to areas with lower or no biocrust cover, possibly due to biocrusts incorporating P into biomass which prevents adsorption onto mineral surfaces (Crain et al., 2018). Additionally, as described in greater detail below, biological and chemical crusts can lead to surface flow of water after precipitation events in our study site and others (Eldridge et al., 2020; Ferrenberg et al., 2017), which can lead to the translocation of organic matter and soil particles away from shrubs to interspaces thereby reducing differences in soil nutrient pools among locations as demonstrated for shrub and interspace patches of *P. glandulosa* in our study system (Parsons et al., 2003). Regardless of the mechanisms underlying the patterns of soil P concentration that we observed, soil P could either be shaping, or responding to plant species composition and cover at a broader spatial scale than soil N or C in our study site, highlighting the need for additional studies of dryland soil biogeochemistry that consider spatial patterns in biotic and abiotic processes.

### 4.2 Solar radiation and soil temperature

The importance of shading and the associated reduction in soil and air temperature for protégé plants beneath the canopy of nurse-plants has been established in numerous ecosystems and for a variety of plant functional types (Castanho & Prado, 2014; Filazzola & Lortie, 2014; Franco & Nobel, 1989; Ren et al., 2008). Both *P. glandulosa* and *L. tridentata* similarly reduced the amount of photosynthetically active radiation (PAR) reaching the surface beneath their canopies, which in turn reduced near-surface soil temperatures that would be experienced by seeds, seedlings, and near-surface roots of co-occurring plants (Figures 4, 5, 9, and S6). This difference in soil temperatures, particularly at peak sunlight hours, has been implicated as a driver of nurse-protégé interactions among shrubs and cacti and among other plant functional types (Reyes & Moya, 2002). A reduction in solar radiation, even when minor, can greatly enhance germination rates and seedling survival in drylands and numerous studies have found increased plant occurrence in areas protected from incident radiation not only by plant canopies, but also by rocks and surface features (Flores & Jurado, 2003; Larmuth & Harvey, 1978).

While shading can be the primary driver of the nurse-plant effect in some situations or systems, the importance of shading as part of a nurse-plant effect is presumably greatest in low-latitude systems or in systems where a substantial portion of plant recruitment and growth occurs while solar radiation is at or near peak annual levels (e.g., summer monsoon dominated systems). As a result, seasonal variation in radiation and temperature could alter not only the degree, but also the dominant driver(s) of the nurse-plant effect. At the same time, plant form and size should influence the level of shading and temperature reduction provided by nurse plants. For example, canopy architecture of *L. tridentata* affects the degree of shading that branches and leaves cause throughout both seasonal and diurnal cycles of solar radiation; variation in shading by different portions of the shrub canopy are least pronounced at times near or during the summer solstice (Neufeld et al., 1988). Thus, the characterization of shading effects in our study—which occurred in the weeks just after the summer solstice—are likely less variable within and among shrubs relative to other times of the year and other dryland systems at higher latitude. How the total nurse-plant effect actually shapes plant population and community dynamics, however, would be further influenced by interannual climate conditions. Specifically, dry years would have less recruitment/growth while wetter years would have more, thus creating scenarios where the realized nurse-plant effect on protégés is smaller and larger, respectively. Additional observational and experimental studies that measure the importance of shading across seasons, latitudes, and years are required if we are to accurately quantify the influence of shading and temperature reduction on dryland plant dynamics.

### 4.3 Soil moisture

At max saturation in lab trials, soil from beneath *P. glandulosa* held a larger volume of water than soil from interspaces and beneath *L. tridentata* likely due to the higher organic matter content of *P. glandulosa* soils (Figure S4). However, sensors deployed in the field revealed no significant difference in VWC beneath the two species and consistently greater soil moisture in unvegetated interspaces (Figure 7). The difference in water volume among soils beneath shrubs versus interspaces was not caused by a different number of soil moisture pulses reaching the surface (Figure 8)—which might be expected in the absence of shrub biomass to intercept or scatter rainfall (Martinez-Meza & Whitford, 1996; X. P. Wang et al., 2011). Surface topography was the likely driver of higher VWC in soils of interspaces versus beneath shrubs; areas beneath shrubs were commonly elevated to a small degree relative to neighboring interspaces leading to surface water flow away from shrubs toward interspaces. Dryland soils are also frequently covered by chemical and/or biological crusts that are temporarily hydrophobic after periods of desiccation and cause rapid ponding and surface runoff in the early stages of precipitation events (Assouline & Mualem, 1997; Eldridge et al., 2020; Thompson et al., 2010). Observations of surface ponding and runoff away from shrubs to interspaces by the authors during this and other studies (Figure S8) further support the hypothesis that local topography as a likely driver of the soil VWC patterns among shrubs and interspaces.

Our observation of higher moisture in interspace soil than under shrubs conflicts with many prior reports of the opposing pattern (e.g., Hong-Fei et al., 2010; Li et al., 2010; Xie et al., 2022), including wetter soils under the shrub *Flourensia cernua* (tarbush) than interspaces in areas neighboring our study site (Kidron & Gutschick, 2013). Many of the available studies of soil moisture in drylands have focused on hillslopes or, conversely, on low lying areas with fine-textured soils while we measured soil moisture of a gently sloping alluvial site with relatively coarse-textured soils. This difference in soil type and landscape position likely has a great influence on soil moisture patterns observed across studies—younger/coarser soils tend to permit greater water infiltration and persistence compared to older/finer soils (Hamerlynck et al., 2002). These same properties can further affect vegetation cover, density, and physical form in ways that feedback to further influence soil moisture near the ground surface (Gearon & Young, 2021; Hamerlynck et al., 2002; Marston, 2010; S. Wang et al., 2021; X. P. Wang et al., 2011).

Beyond variation in surface topography and soil permeability among studies, measurements of soil moisture are significantly affected by sampling frequency and depth in the soil column. Differences among the soil water contents we observed for shrub versus interspace were derived from a time series of 15-minute intervals while values reported by Kidron & Gutschick (2013), for example, were collected once to only a few times per week. Additionally, we focused on near-surface water content because of its importance for seed moisture and germination and seedling establishment while many prior studies measured soil water content at depths of 20 cm or greater where recent germinants and annual plants are less likely to be rooted. Future efforts to measure surface flow and soil moisture content across depth and in relation to variation in surface topography, as well as shrub size and architecture, across different climate types could greatly advance our understanding of spatiotemporal variation in soil moisture availability and its relative influence on spatial patterns of plant occurrence in dryland systems.

### 4.4 Nurse plant effects and shrub size

The canopy size of *L. tridentata* and *P. glandulosa* had a significant influence on understory shading, while canopy size of *P. glandulosa* was also related to soil biogeochemistry. Whether this latter result was caused by or simply correlated with shrub canopy size, however, was unclear as rooting properties and other attributes likely vary in relation to canopy cover. Given the effects of shrub size measured here and elsewhere, advances in understanding nurse plant dynamics could come from a greater consideration of variation in shrub form and size—attributes that change between species, across time as shrubs grow and acclimate to local conditions, and across gradients in resources, climate, topography, and geomorphology (Gearon & Young, 2021; Hamerlynck et al., 2002). Not surprisingly, the importance of the nurse-plant effect and its drivers can change across these same gradients (Cavieres et al., 2006; Tewksbury & Lloyd, 2001; Tielborger & Kadmon, 2000). Further investigation of how variation in shrub size and form affect interactions with emerging and understory plants has also gained importance in light of increasing rates of disturbance in drylands. As global change pressures continue to intensify in drylands (Prudhomme et al., 2014; Sloat et al., 2018), a potential loss of nurse plant cover and reduction in size as disturbances cause shrub mortality or dieback could have substantial effects on soil biogeochemical pools and processes in addition to driving losses in plant species richness and diversity.

### 4.5 Conclusions

Dryland ecosystems are characterized by chronic and/or pulsed aridity that shape the distribution and abundance of plant species. Perennial plants can modify soil and microclimatic conditions beneath their canopies, creating spatial variation in the level and types of environmental stresses in ways that can facilitate recruitment and survival of co-occurring plants. Many mechanisms can produce plant aggregations. When and where the difference mechanisms are most important and how they interact over resource/climate gradients remains largely unclear across dryland ecosystems (Filazzola & Lortie, 2014; Flores & Jurado, 2003). While it is commonly assumed that higher soil moisture under perennial shrubs is the primary driver of the ‘nurse-plant effect’ in warm desert systems, we found that unvegetated interspaces consistently had a larger volume of soil water near the ground surface than areas beneath shrubs. Nevertheless, the higher levels of solar radiation and concomitant soil temperatures in interspaces likely lead to greater physiological stress in seeds and plants as they emerge from the seed bank, particularly in early life stages when root systems are either too shallow or limited to access deeper soil water. Once rooted, plants could subsequently benefit from the larger concentration of nitrogen observed in soils collected from beneath shrub canopies; globally, higher leaf nitrogen is associated with greater photosynthetic rates by plants and plant allocate a greater proportion of leaf nitrogen to the photosynthetic enzyme ribulose-1,5-bisphosphate carboxylase-oxygenase (RuBisCO) as aridity increases (Luo et al., 2021). Further understanding of the mechanisms that lead to plant aggregation will require experimental efforts that aim to manipulate specific pathways across different plant communities and clines in resources and climate.

## Acknowledgements

This work was supported by a USDA-NIFA-AFRI (#2019-67020-29320) award to S.F. This work is also in thanks to support from the L’Oréal USA For Women in Science Fellowship and American Association for the Advancement of Science (AAAS) awards to B.O. We thank Sasha Reed and Robin Reibold (USGS, Moab, UT) for support with biogeochemical analyses, Jon Anderson (New Mexico State University/Jornada Basin LTER) and Andrew Cox (New Mexico State University’s Chihuahuan Desert Rangeland Research Center) for logistical support and Megan Rabinowich, Andrew Dominguez, and Matthew Tryc for assistance with data collection. Any use of trade, product or firm names is for descriptive purposes only and does not imply endorsement by the U.S. Government.

## Supplemental Materials

**Table S1.** Mean values^†^ of shrub density, shrub height, and distance to nearest shrub neighbor in study blocks where creosote bush (*L. tridentata*) or honey mesquite (*P. glandulosa*) are the dominant cover.

| Species | Density | (± 1SE) | Height | (± 1SE) | Nearest | (± 1SE) |
| --- | --- | --- | --- | --- | --- | --- |
| *L. tridentata* | 2.13 | 0.23 | 83.88 | 7.06 | 129.60 | 7.69 |
| *P. glandulosa* | 5.70 | 0.32 | 87.70 | 2.39 | 73.33 | 5.73 |

<sup>†</sup>Values are calculated from all shrub species and not limited to conspecifics.

**Table S2.**
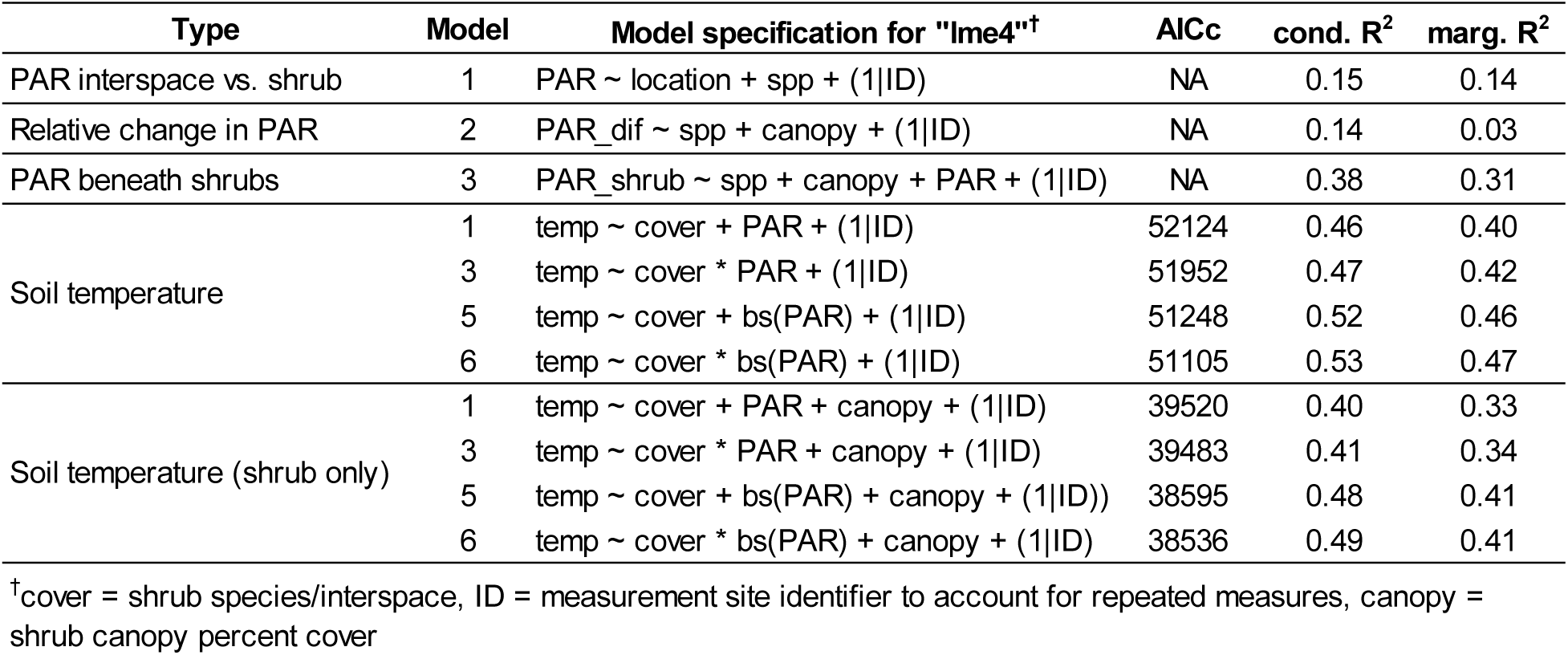
Candidate models for PAR and soil temperature.

**Table S3.** VWC corrections for ECHO-EC5 sensors in the field setting.

| Provenance | Laboratory-derived correction model |
| --- | --- |
| Interspace | $\text{Corrected\_vwc} = 0.5915773 * \text{VWC} + 0.033967 * \text{Location} + -0.073358$ |
| *L. tridentata* | $\text{Corrected\_vwc} = 0.623161 * \text{VWC} + 0.009321 * \text{Location} + -0.060295$ |
| *P. glandulosa* | $\text{Corrected\_vwc} = 0.736250 * \text{VWC} + -0.005914 * \text{Location} + -0.063372$ |

**Table S4.** MANOVA table for the effects of shrub species and location (interspace vs. under shrub) on soil biogeochemical pools.

| **Factor** | **Df** | **Pillai** | **Approx. F** | **P** |
| --- | --- | --- | --- | --- |
| Intercept | 1 | 0.968 | 188.501 | < 0.0001 |
| Shrub spp. | 1 | 0.308 | 2.787 | 0.0124 |
| Location | 1 | 0.496 | 6.154 | < 0.0001 |
| Residuals | 57 |  |  |  |

**Table S5.** Mixed model results assessing the influence of shrub species, canopy cover, and total incoming PAR measured in open interspaces on PAR measured at the ground-level under shrub canopies.

| **Factor** | **Estimate** | **SE** | **df** | **t** | **P** |
| --- | --- | --- | --- | --- | --- |
| Intercept | 259.40 | 133.10 | 27.03 | 1.95 | 0.0617 |
| Shrub species | 16.37 | 43.97 | 27.02 | 0.37 | 0.7125 |
| Canopy cover | -451.10 | 212.10 | 27.01 | -2.13 | 0.0427 |
| Incoming PAR | 0.31 | 0.00 | 55180.00 | 159.29 | < 0.0001 |

**Table S6.**
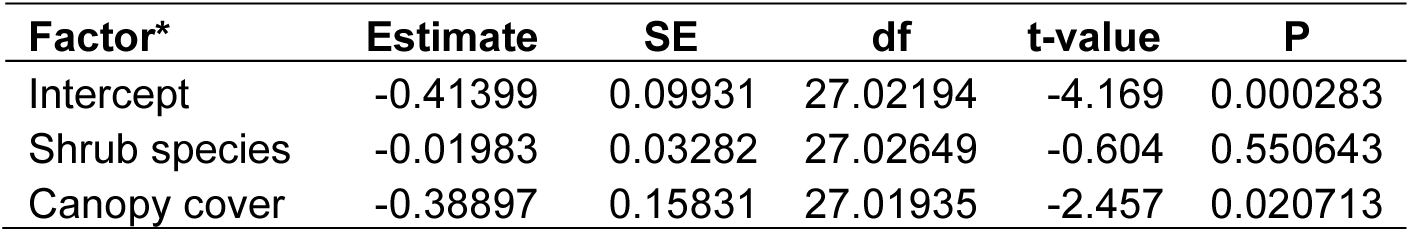
Best-fit linear mixed model of the relative reduction in incoming PAR beneath shrub canopies.

| **Factor\*** | **Estimate** | **SE** | **df** | **t-value** | **P** |
| --- | --- | --- | --- | --- | --- |
| Intercept | -0.41399 | 0.09931 | 27.02194 | -4.169 | 0.000283 |
| Shrub species | -0.01983 | 0.03282 | 27.02649 | -0.604 | 0.550643 |
| Canopy cover | -0.38897 | 0.15831 | 27.01935 | -2.457 | 0.020713 |

**Table S7.**
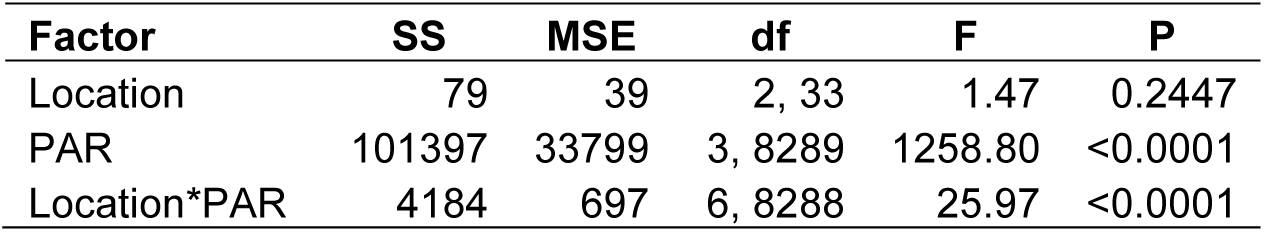
Best fit mixed-model result for soil temperature as a function of incoming photosynthetically active radiation (PAR) and location (open-interspace or beneath the canopy of *L. tridentata* or *P. glandulosa* shrubs)

**Table S8.** Mixed-model results relating soil VWC and proportional soil saturation to soil collection location, cover type, and time since reaching saturation.

| Model | Factor | Estimate | S.E. | d.f. | t | P |
| --- | --- | --- | --- | --- | --- | --- |
| Volumetric water content | Intercept | 0.20770 | 0.0059 | 34.2 | 35.0 | < 0.0001 |
|  | *L. tridentata* soil | 0.01287 | 0.0072 | 32.5 | 1.8 | 0.0827 |
|  | *P. glandulosa* soil | 0.05484 | 0.0072 | 32.3 | 7.6 | < 0.0001 |
|  | Time (minutes) | -0.00001 | 0.0000 | 489.7 | -70.2 | < 0.0001 |
|  | Site | 0.02272 | 0.0059 | 32.3 | 3.9 | 0.0005 |
| Proportion max saturation | Intercept | 0.81530 | 0.0265 | 32.9 | 30.8 | < 0.0001 |
|  | *L. tridentata* soil | 0.07885 | 0.0322 | 32.2 | 2.4 | 0.0200 |
|  | *P. glandulosa* soil | 0.08777 | 0.0322 | 32.1 | 2.7 | 0.0103 |
|  | Time (minutes) | -0.00005 | 0.0000 | 487.2 | -83.0 | < 0.0001 |
|  | Site | 0.02541 | 0.0263 | 32.0 | 1.0 | 0.3406 |

**Figure S1.**
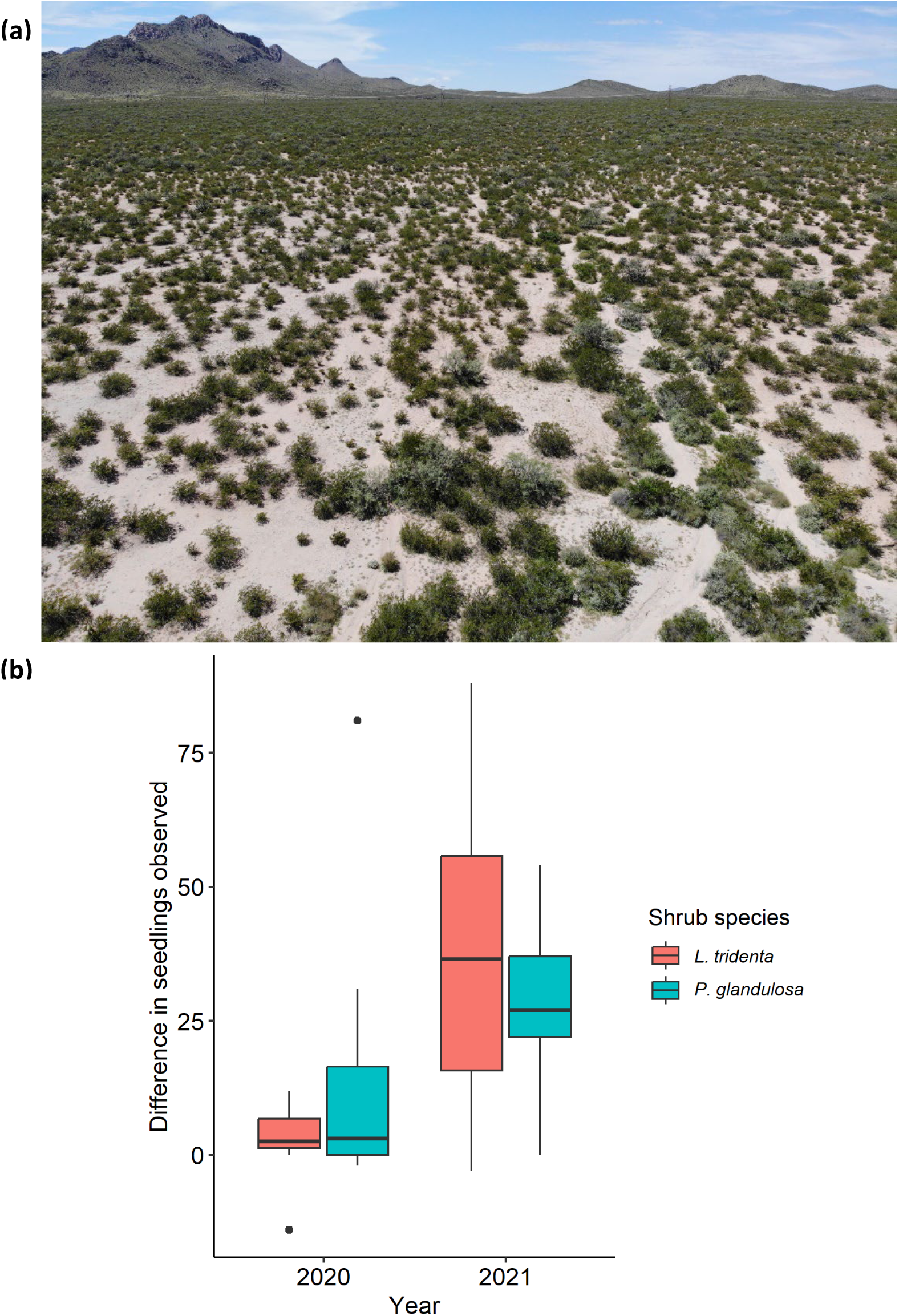
**(a)** Aerial view of the western portion of the study site located on the Chihuahuan Desert Rangeland Research Center bordering the Jornada Basin Experimental Range in southern New Mexico, USA. **(b)** Difference in seedlings/plants observed (Δ = shrub – interspace) in paired plots under shrubs and interspaces for 2020 (below average monsoon rainfall) and 2021 (above average monsoon rainfall).

**Figure S2.**
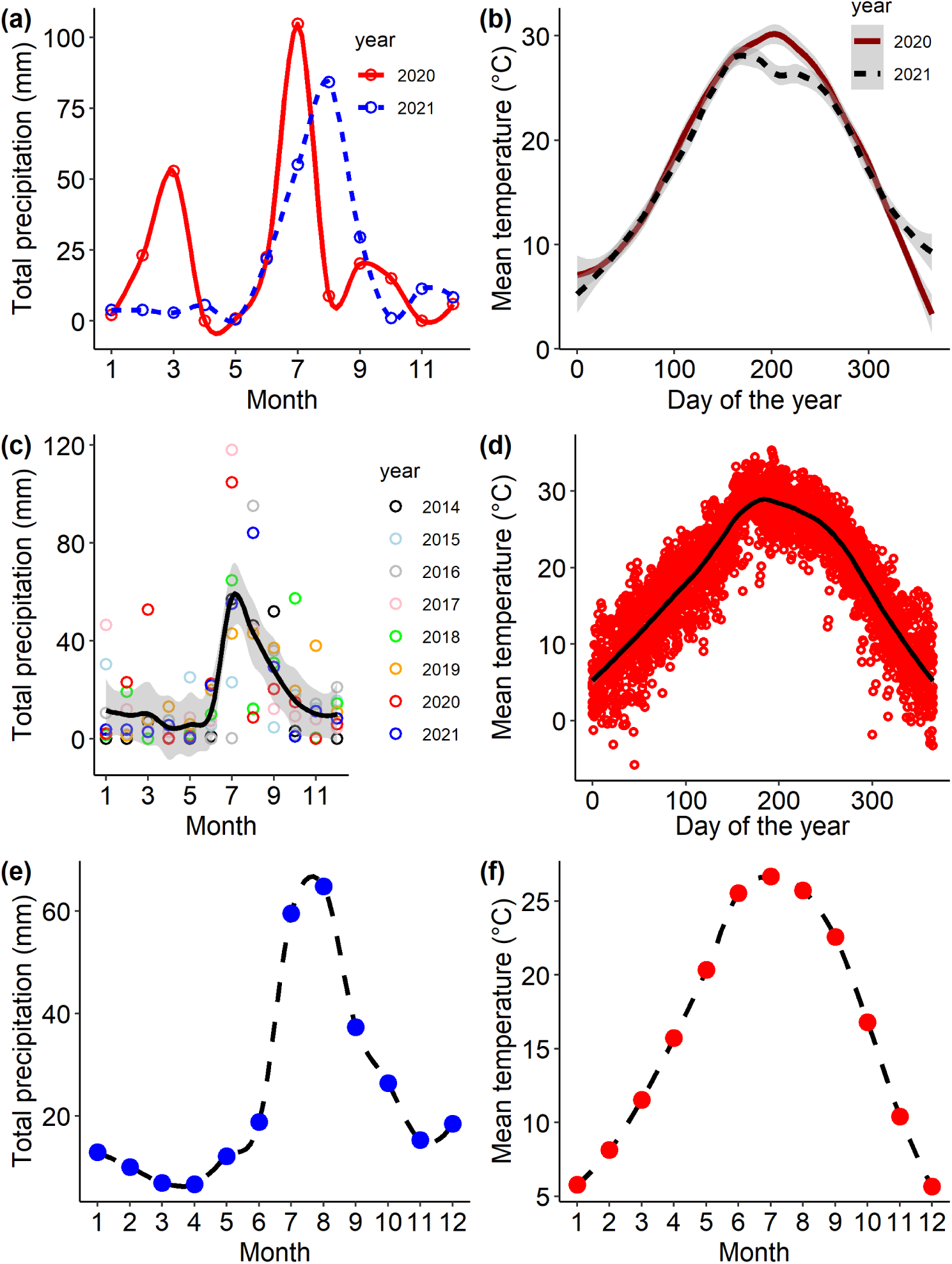
Monthly climate values for 2020 through 2021(panels a and b) and monthly and daily climate values (panels c – f) recorded from September 2013 through December 2021 at a climate station roughly 1 km from the described study site.

**Figure S3.**
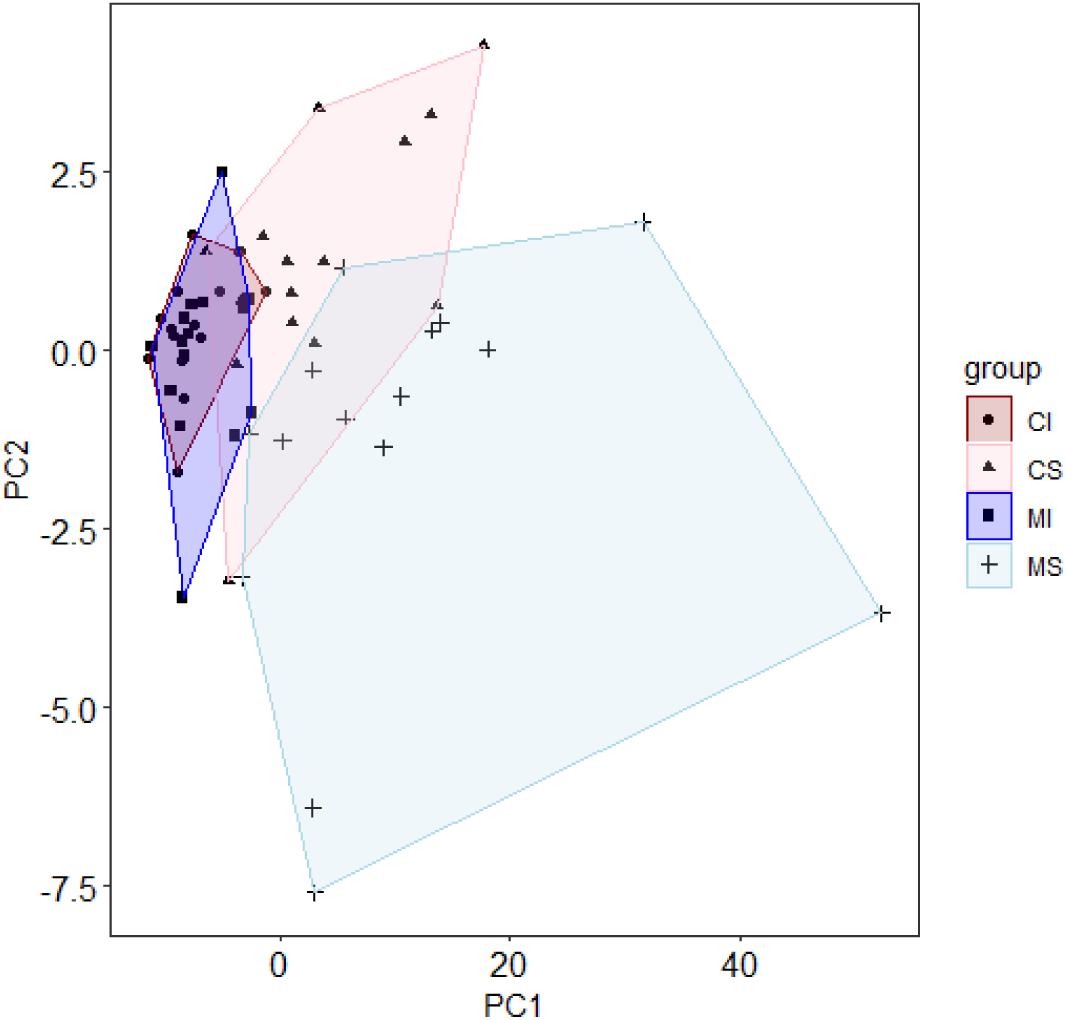
Local Fisher discriminant analysis (LFDA) plot illustrating the multivariate biogeochemical pools in soil samples collected beneath shrubs of *L. tridentata* (creosote, CS) and *P. glandulosa* (mesquite, MS) and in neighboring, unvegetated interspaces (CI and MI) that were paired with each shrub.

**Figure S4.**
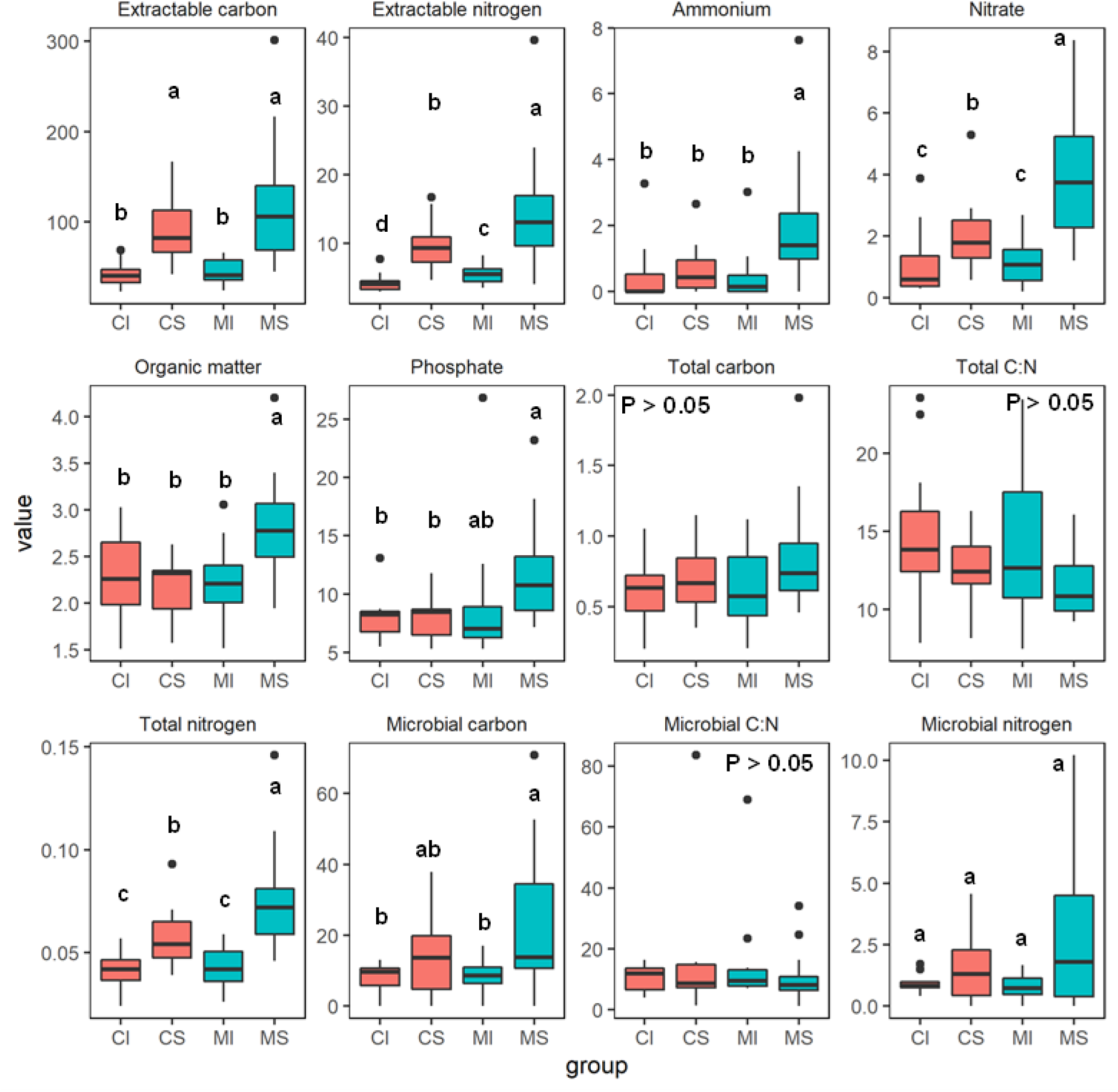
Biogeochemical pools of the top 10 cm of soil collected from interspaces and beneath shrubs in locations with L. tridentata (creosote) and P. glandulosa (mesquite). “C” and “M” on the x-axis refer to “creosote” and “mesquite” while “I” and “S” denote whether the sample was from an “interspace” neighboring the species or collected from beneath a “shrub” of the species.

**Figure S5.**
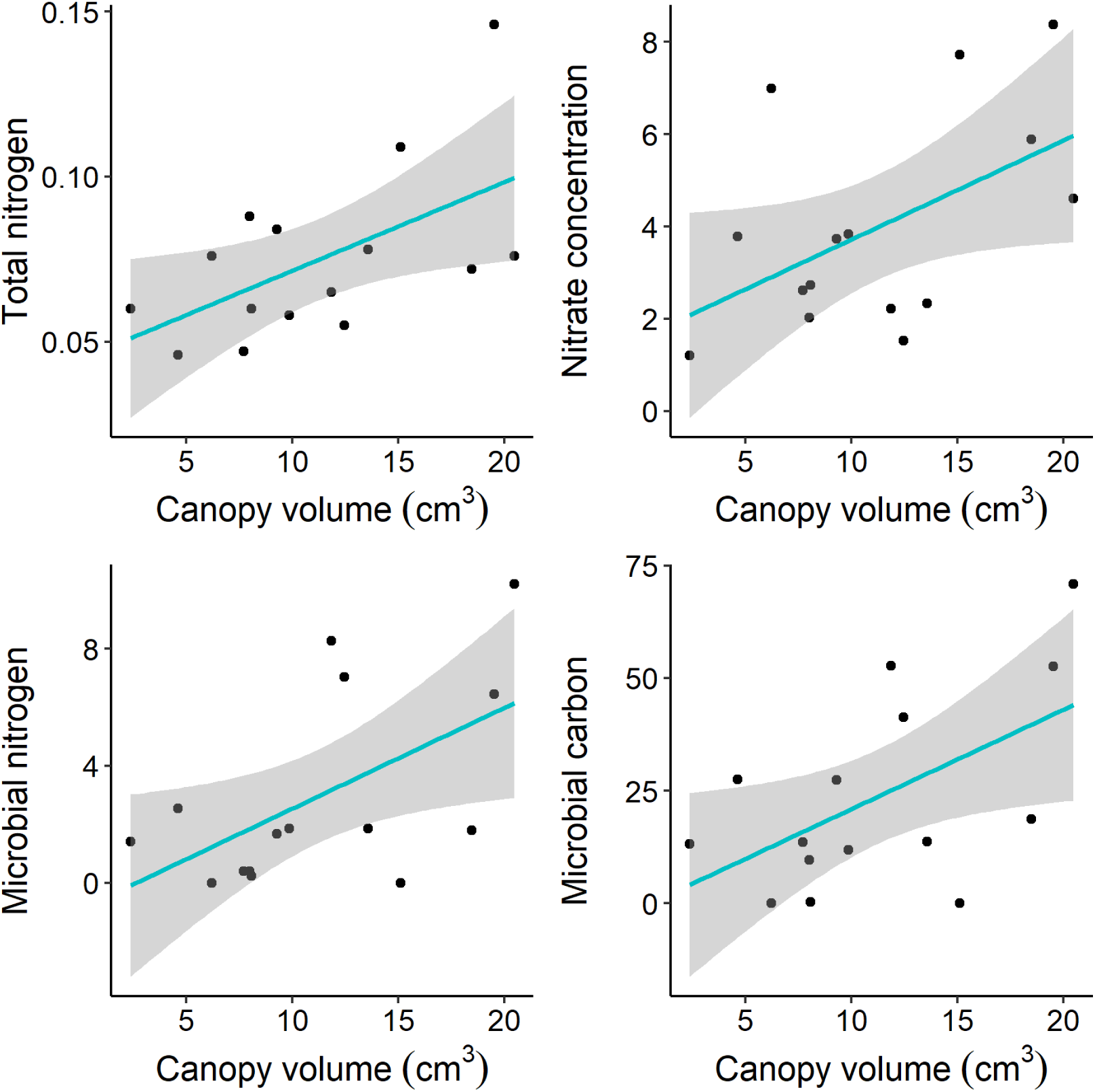
Influence of the of size, calculated as canopy volume, of *Prosopis glandulosa* shrubs on biogeochemical pools.

**Figure S6.**
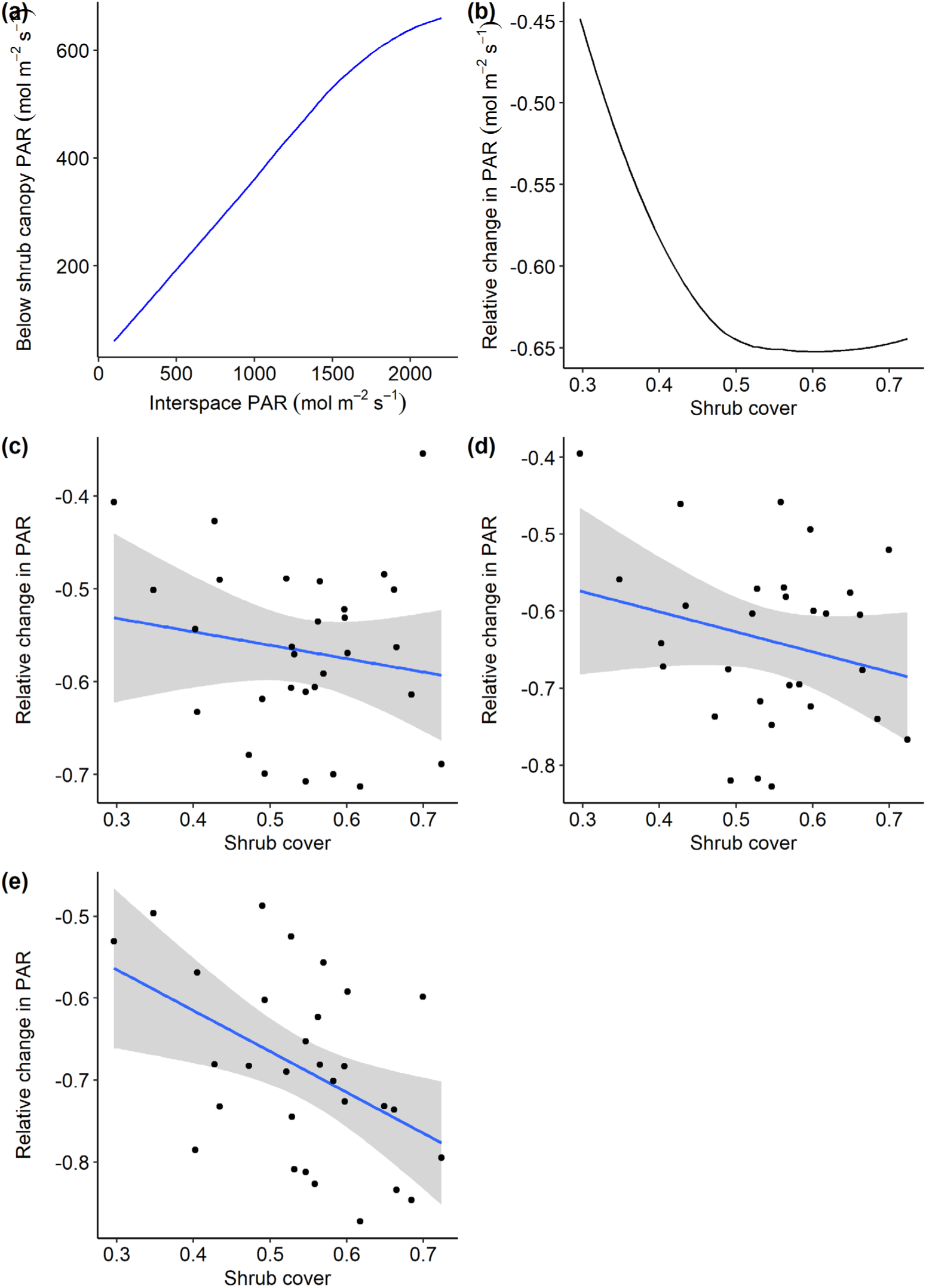
(a) Relationship between photosynthetically active radiation (PAR, mol m^-2^ s^-1^) observed in open interspaces vs. under shrub canopies, (b) the relative change in PAR as a function of proportional shrub-canopy cover (from upward facing photographs)—the lines fit in (a) and (b) represent the population mean value derived from repeated measures within multiple shrubs. Remaining panels show the mean reduction in PAR by shrub proportional cover at increasing levels of incoming PAR from (c) “low” (100 – 800 mol m^-2^ s^-1^), to (d) “moderate” (801 – 1500 mol m^-2^ s^-1^), and (e) “high” (1501 – 2200 mol m^-2^ s^-1^).

**Figure S7.**
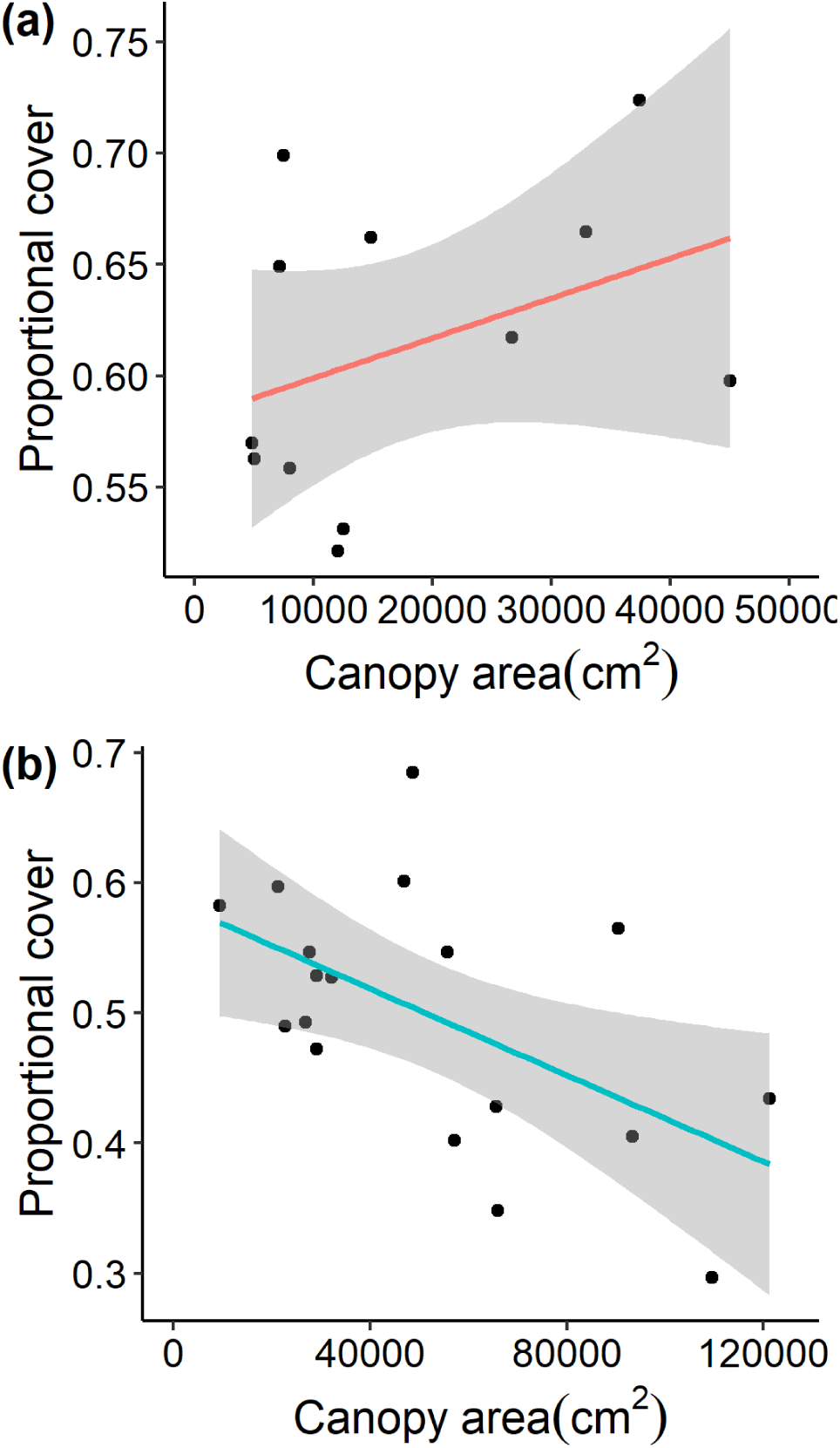
The relationship between proportional cover (area in cm^2^ covered by shrub limbs and leaves in upward facing photos) and shrub canopy area calculated as an oval based on the longest set of perpendicular axes for (a) *L. tridentata* (creosote; R^2^ = 0.14, P = 0.228) and (b) *P. glandulosa* (mesquite; R^2^ = 0.30, P = 0.018).

**Figure S8.**
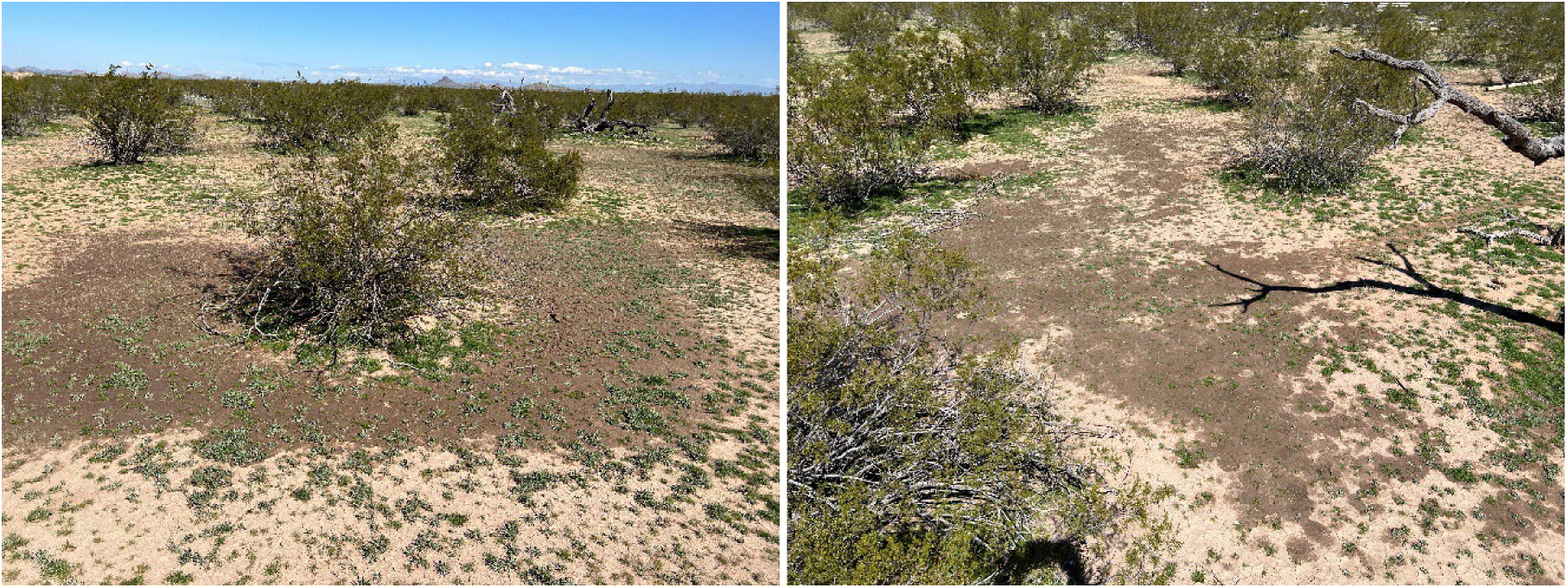
Photos illustrating a higher level of near-surface soil moisture away from the canopies and in low-lying interspaces around *L. tridentata* shrubs following a recent rain event in the Sonoran Desert of the U.S. Southwest (photos by Armin Howell).

